# Dual inhibition of GTP-bound (ON) and GDP-bound (OFF) KRAS^G12C^ suppresses PI3Kα and leads to potent tumor inhibition

**DOI:** 10.64898/2026.04.27.718135

**Authors:** Katherine Parker, Samar Ghorbanpoor, Wafa Malik, Lauren Highfield, Jamie Wang, Emily Hensley, Grace Kelley, Sarah Clark, Michela Ranieri, Soumyadip Sahu, Cathy Zhang, Magdalena Ploszaj, Daniel Andrussier, Hsin-Yi Huang, Ting Chen, Bin Wang, Rui Xu, Saman Setoodeh, Ken Lin, Anna E. Maciag, Frank McCormick, Pedro J. Beltran, Kwok-Kin Wong, James P. Stice, Kerstin W. Sinkevicius, Aaron N. Hata

## Abstract

Current approved KRAS^G12C^ inhibitors covalently bind the inactive GDP-bound (OFF) form of KRAS^G12C^. Recently, KRAS^G12C^ inhibitors that selectively bind to the GTP-bound (ON) form of both KRAS^G12C^ (ON) and (OFF) forms have been reported and entered clinical testing. In principle, KRAS^G12C^ (ON) inhibitors may be less susceptible to adaptive mechanisms that promote resistance to (OFF) inhibitors, however the specific mechanisms that differentiate the activity of (ON) versus (OFF) inhibition are not well understood. We profiled the activity of BBO-8520, a covalent dual inhibitor of GTP-bound (ON) and GDP-bound (OFF) KRAS^G12C^, in *KRAS^G12C^*-mutant non-small cell lung cancer models. BBO-8520 exerted more potent and sustained inhibition of KRAS^G12C^ and anti-tumor activity *in vitro* and *in vivo* compared with sotorasib, a KRAS^G12C^ (OFF)-only inhibitor. While cells treated with BBO-8520 or sotorasib both exhibited feedback reactivation of MAPK signaling driven by wild-type HRAS/NRAS isoforms, more durable suppression of KRAS^G12C^ by BBO-8520 was associated with decreased PI3Kα-AKT activation *in vitro*. Disruption of the interaction between RAS and PI3Kα using a novel protein:protein interaction inhibitor suppressed PI3Kα-AKT activation and increased the tumor response to sotorasib to a similar level as BBO-8520. Moreover, in some contexts, disruption of RAS-PI3Kα further increased the anti-tumor activity of BBO-8520 monotherapy. These results reveal mechanistic differences between KRAS (ON) and (OFF) inhibitors, highlight the importance of PI3Kα-AKT signaling in driving resistance to KRAS inhibition in lung cancer, and suggest combination strategies that suppress PI3Kα-AKT to improve the response to KRAS inhibitors.

## Introduction

*RAS* is one of the most frequently mutated oncogenes in cancer, with mutations in the KRAS isoform commonly found in non-small cell lung cancer (NSCLC), colorectal cancer (CRC) and pancreatic ductal adenocarcinoma (PDAC)^1,2^. RAS proteins are small GTPases, which cycle between the GTP-bound (ON) and GDP-bound (OFF) conformations^3,4^. Oncogenic *KRAS* mutations disrupt the interaction between KRAS and GTPase activating proteins (GAPs), which facilitate the hydrolysis of GTP and convert KRAS from ON to OFF, thus stabilizing KRAS in its GTP-bound active conformation^5^. Constitutively active GTP-bound KRAS activates downstream mitogen-activated protein kinase (MAPK) and phosphatidylinositol 3-kinase (PI3K) pathways, which promote cell survival, proliferation, and differentiation^3,4^.

First-generation KRAS inhibitors (sotorasib, adagrasib), which selectively target the KRAS^G12C^ mutation, have been approved for treatment of KRAS^G12C^-mutant NSCLC and CRC (in combination with EGFR-blocking antibodies)^6–8^. These agents inhibit the KRAS^G12C^-mutant protein by covalently binding to the mutant cysteine in the GDP-bound conformation, locking KRAS in the OFF state^9,10^. Due to GAP-independent, intrinsic hydrolysis of GTP bound to KRAS^G12C^, KRAS inhibition is achieved as the cellular pool of KRAS^G12C^ is cumulatively trapped in the inactive, GDP-bound conformation^11^, depleting the pool of GTP-bound (ON) KRAS available to engage with effectors.

Despite the success of first-generation KRAS^G12C^ inhibitors, partial responses are induced in only 30-40% of NSCLC patients and durable responses are rare^6,7^. Preclinical studies have demonstrated that these agents are susceptible to adaptive reactivation of MAPK signaling, often driven by receptor tyrosine kinases (RTKs)^12–14^. Specifically, high levels of basal or adaptive RTK signaling can increase levels of the GTP-bound (ON) conformation of KRAS^G12C^ and diminish the availability of the GDP-bound (OFF) conformation targeted by first generation inhibitors. Moreover, the high intracellular concentration of GTP^15^ likely ensures that newly synthesized KRAS^G12C^ protein initially enters the GTP-bound (ON) pool. These data suggest that inhibiting the GTP-bound (ON) state may more rapidly and durably suppress KRAS^G12C^ and lead to deeper and more sustained efficacy. On the other hand, other preclinical studies have suggested that MAPK reactivation is mediated by non-mutant RAS isoforms such as HRAS and NRAS^14,16,17^ that would not be susceptible to improved targeting of KRAS^G12C^. Thus, it remains an open question whether targeting GTP-bound (ON) KRAS versus GDP-bound (OFF) KRAS represents a superior approach.

BBO-8520, a first-in-class direct and covalent dual inhibitor of GTP-bound (ON) and GDP-bound (OFF) KRAS^G12C^ is currently under clinical development for *KRAS^G12C^*-mutant NSCLC (NCT06343402). In a recently reported interim analysis, BBO-8520 demonstrated an objective response rate (ORR) of 65% (11/17) and a 6-month progression-free survival (PFS) rate of 66% in patients with KRAS^G12C^-mutant NSCLC across all dose levels, with a generally well-tolerated and manageable safety profile^18^. In preclinical studies, BBO-8520 exhibited more rapid target engagement and improved potency and efficacy compared to first-generation KRAS^G12C^ inhibitors in *KRAS^G12C^*-mutant cell lines and xenograft models^19^. Here, we investigated the mechanistic basis for the improved efficacy of dual (ON) and (OFF) targeting. We find that BBO-8520 achieves more durable suppression of KRAS^G12C^ compared to sotorasib, yet comparable suppression of downstream MAPK signaling. In contrast, BBO-8520 suppressed PI3K-AKT to a greater extent than sotorasib, and equivalent to sotorasib combined with BBO-10203, a covalent inhibitor of the interaction between RAS and PI3K p110α^20^. Moreover, the combination of sotorasib and BBO-10203 achieved comparable anti-tumor activity in vitro and in vivo, highlighting the importance of suppressing PI3K-AKT signaling therapeutic response to KRAS inhibition in NSCLC. These results provide mechanistic insights into the differences between dual KRAS^G12C^ (ON) and (OFF) versus (OFF) inhibitors and provide rationale for combining inhibitors of RAS-PI3K/p110α with KRAS inhibitors.

## Results

### BBO-8520 exhibits potent activity against *KRAS^G12C^*-mutant NSCLC cells

To broadly assess the potential benefit of dual targeting GTP-bound (ON) and GDP-bound (OFF) KRAS^G12C^, we compared the activity of BBO-8520^19^ and related tool compound BBO-7410 to the GDP-bound (OFF) inhibitor sotorasib^1^ across a panel of 14 *KRAS^G12C^*-mutant NSCLC cell lines with diverse co-occurring mutations (Supplemental Figure 1A). Like prior studies of KRAS^G12C^ OFF inhibitors^9,10^, we observed wide variability in sensitivity to sotorasib across the cell line panel in short term proliferation assays, with IC_50_ and E_max_ values ranging from 6-350 nM and 16-95%, respectively (Figure 1A-E, Supplemental Figure 1B). Sensitivity to BBO-8520 (E_max_ range 18-91%) and related compounds was also variable across the cohort and correlated with sotorasib sensitivity (Supplemental Figure 1B, C), but comparable maximal effects on cell proliferation were consistently achieved at ∼100-fold lower drug concentrations (BBO-8520 IC_50_ 0.05-1.5 nM; Figure 1C) and ∼375-fold lower free drug concentrations after accounting for the fraction unbound in 10% FBS-containing media^19^. Consistent with this, sotorasib suppressed downstream phospho-ERK and phospho-RSK signaling at concentrations above 10-100 nM, while BBO-8520 did so at doses as low as 1 nM (Figure 1F, Supplemental Figure 1D). Although this dramatic increase in potency did not translate into increased maximum efficacy when considering the cell line cohort as a group, BBO-8520 exhibited modestly greater E_max_ in 3 cell lines (for example, MGH1062-1; Figure 1A, D) and numerically greater E_max_ values (albeit not statistically significant) were observed in 10 out of 14 of cell lines. No differences in drug sensitivity were observed when stratifying cells according to the presence of co-occurring mutations in TP53 or STK11 (Supplemental Figure 1E).

**Figure 1.**
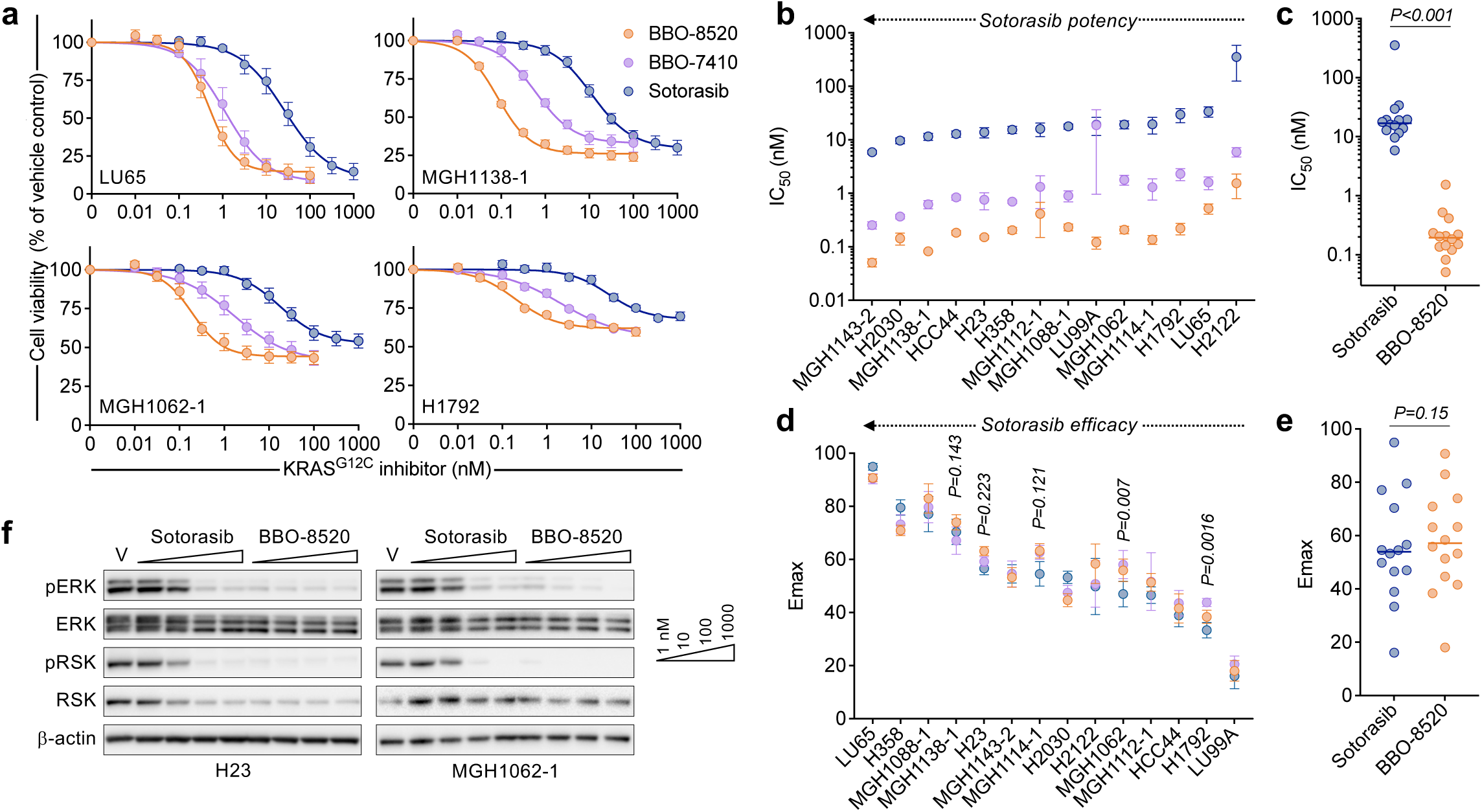
The KRAS^G12C^ (ON) inhibitor BBO-8520 exhibits more potent inhibition of MAPK signaling and cell proliferation compared to the KRAS^G12C^ (OFF) inhibitor sotorasib. a, Composite dose response curves of *KRAS^G12C^*-mutant NSCLC cell lines. Cells were treated with KRAS inhibitors for 72 hours and cell viability was quantified by CellTiter-Glo. Curves shown are mean and S.E.M. of n=3-7 independent biological replicates. b-c, Comparison of IC50 values from 3-day viability assays. Values shown are mean and S.E.M. of n=3-7 independent biological replicates. d-e, Comparison of Emax values from 3-day viability assays. Values shown are mean and S.E.M. of n=3-7 biological replicates. f, Western blot analysis of *KRAS^G12C^*-mutant NSCLC cell lines treated with increasing concentrations of sotorasib or BBO-8520 for 6 hours. Data is representative of n=3 independent biological replicates.

The efficacy of KRAS^G12C^ (OFF) inhibitors is limited by adaptive resistance driven by feedback reactivation of the MAPK pathway, which can be mediated by newly synthesized KRAS^G12C^ that adopts a GTP-bound conformation^12^ or by activation of wild-type RAS isoforms^14,17^. In principle, the former but not the latter, could be overcome by KRAS^G12C^ (ON) inhibition. To assess the activity of BBO-8520 over longer time frames of adaptive resistance, we labeled a sub-cohort of 7 cell lines representing a range of sensitivities with GFP to enable live-cell tracking (Supplemental Figure 2A). After confirming that the response to KRAS^G12C^ inhibitors remained consistent after GFP labelling (Supplemental Figure 2B), we treated the cells with 10 nM, 100 nM or 1 μM BBO-8520 or 1 μM sotorasib and monitored cell numbers for up to 20 days (Figure 2A, Supplemental Figure 2C). In contrast to the short-term results, these longer-term assays revealed that BBO-8520 suppressed cell proliferation to a greater extent than sotorasib, even at doses as low as 10 nM (Figure 2B, Supplemental Figure 2D). To account for differences in growth rates and KRAS^G12C^ inhibitor sensitivities, we chose a representative timepoint during the 5 – 10 days after initiation of treatment for quantification of drug effect. In all 7 cell lines surveyed, we observed a significantly greater decrease in cell proliferation with 100 nM (20 nM free) BBO-8520 compared with 1 μM (750 nM free) sotorasib, and in 4 of 7 cell lines treated with 10 nM (2 nM free) BBO-8520 (Figure 2C-D, Supplemental Figure 2E). These data indicate that potent inhibition of both the GTP-bound (ON) and GDP-bound (OFF) conformations of KRAS^G12C^ leads to greater suppression of cell proliferation over time.

**Figure 2.**
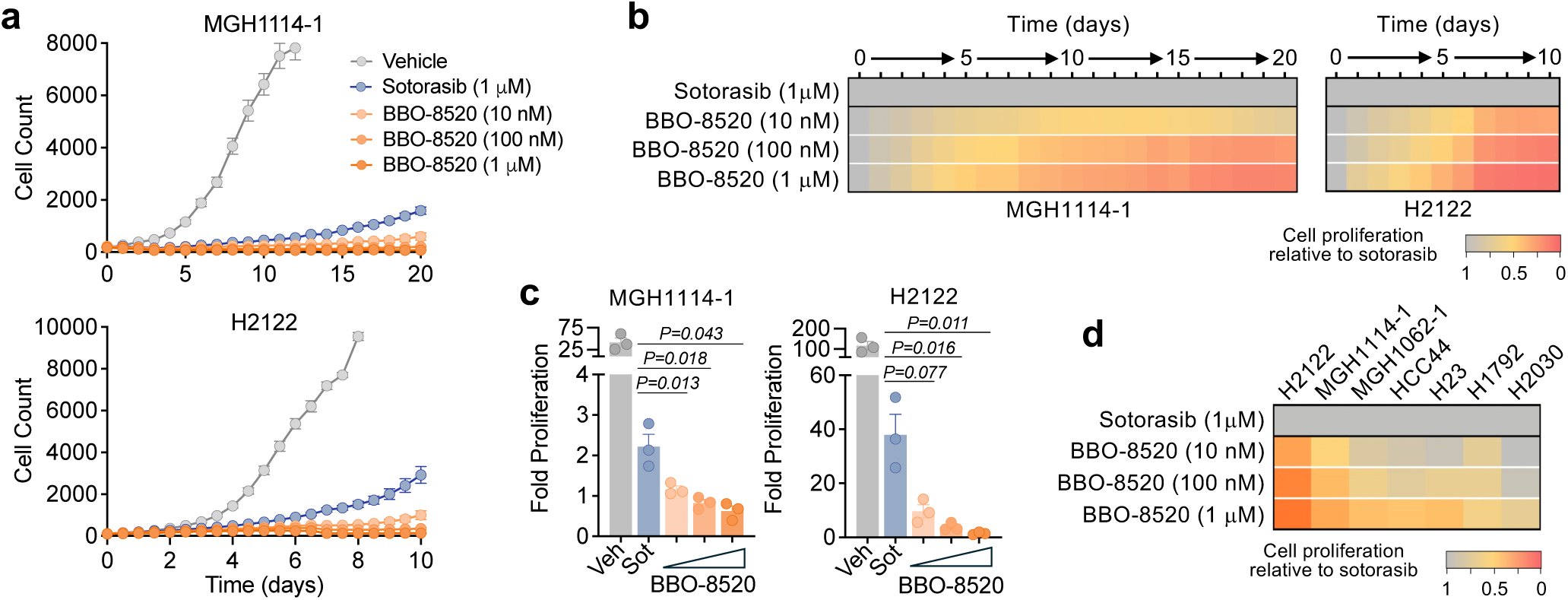
BBO-8520 exhibits more durable suppression of cell proliferation compared to sotorasib. **a,** Imaged-based monitoring (Incucyte) of proliferation of *KRAS^G12C^*-mutant NSCLC cell lines treated with sotorasib or BBO-8520. Data are mean and S.E.M of n=4-6 technical replicates and are representative of n=3 independent biological replicates. **b,** Relative cell proliferation of cells treated with BBO-8520, normalized to sotorasib. Values are mean of n=3 biological replicates, corresponding to the cell counts shown in panel A. **c,** Comparison of cell proliferation after treatment with 10 nM, 100 nM or 1 μM BBO-8520 or 1 μM sotorasib. Values represent cell proliferation relative to baseline determined after 10 days from three independent biological replicates. **d,** Relative cell proliferation of cells treated with BBO-8520, normalized to sotorasib after 10 days (H2122, MGH1114-1, MGH1062-1, H23, H2030), 6 days (H1792), or 5 days (HCC44) to account for variability in baseline (vehicle-treated) growth rates. Values are mean of n=3-4 biological replicates.

### BBO-8520 achieves durable suppression of KRAS^G12C^ and downstream PI3K-AKT signaling

Given that the differences between BBO-8520 and sotorasib efficacy became more apparent over time, we hypothesized that more durable suppression of KRAS^G12C^ by BBO-8520 might suppress adaptive pathway reactivation. To assess this, we first assessed the level of signaling-competent GTP-bound KRAS^G12C^ in short term assays. We treated cells with 100 nM BBO-8520 or 1 μM sotorasib for 6 or 48 hours followed by RAF-RBD pulldown of RAS-GTP. At these short-term timepoints, BBO-8520 suppressed KRAS^G12C^ engagement with RAF to a greater extent than sotorasib (p=0.009, 6h; p=0.026, 48h) (Figure 3A-B, Supplemental Figure 3A). Next, we treated cells with sotorasib or BBO-8520 for 14 days prior to RAS-GTP pulldown. Compared to vehicle controls, both sotorasib and BBO-8520 treated cells had suppressed engagement levels of KRAS-GTP (Figure 3C-E). However, long-term treatment with BBO-8520 consistently suppressed KRAS-GTP levels to a greater extent than sotorasib (p=0.006).

**Figure 3.**
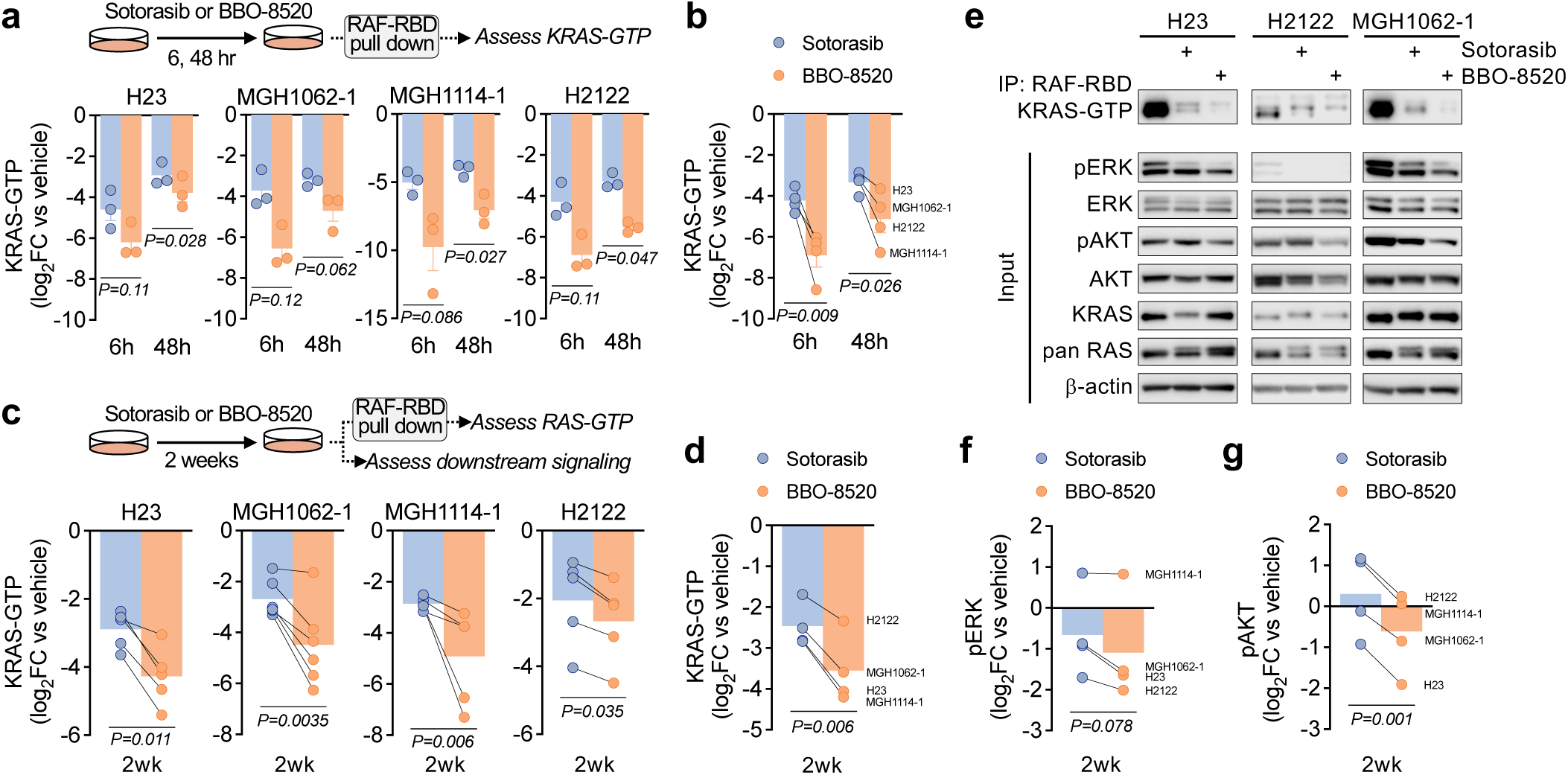
BBO-8520 achieves more durable suppression of KRAS^G12C^-RAF engagement and inhibition of PI3K-AKT compared to sotorasib. **a,** *KRAS^G12C^*-mutant NSCLC cells were treated with BBO-8520 (100 nM) or sotorasib (1 μM) for 6 or 48 hours and RAF-RBD pull-down was performed to assess engagement between KRAS-GTP and RAF. Data are quantified band intensities from western blots (see Supplemental Figure 3a) normalized to vehicle control, n=3 independent biological replicates. **b,** Average change in KRAS-GTP levels after 6 or 48 hours determined by RAF-RBD pull-down. Each data point represents the average change in KRAS-GTP in an individual cell line from panel A. **c,** *KRAS^G12C^*-mutant NSCLC cells were treated with BBO-8520 (100 nM) or sotorasib (1 μM) for 2 weeks and RAF-RBD pull-down was performed to assess engagement between KRAS-GTP and RAF in residual tumor cells. Data are quantified band intensities from western blots normalized to vehicle control, n=4-6 independent biological replicates. **d,** Average change in KRAS-GTP levels after 2 weeks as determined by RAF-RBD pull-down. Each data point represents the average change in KRAS-GTP in an individual cell line from panel D. **e,** Representative western blot images from RAF-RBD pull-down after 2 weeks treatment with sotorasib or BBO-8520, as quantified in panels C and D. **f-g,** Average pERK (F) and pAKT (G) levels after treatment with BBO-8520 (100 nM) or sotorasib (1 μM) determined by western blot band quantification (see panel E). Each dot represents the mean of an individual cell line, n=4-6 independent biological replicates.

Next, we examined whether the more durable suppression of KRAS-GTP led to more durable suppression of downstream signaling. After 14 days of treatment, there was a trend toward decreased phospho-ERK levels in cells treated with BBO-8520 compared to sotorasib, however this did not reach statistical significance (p=0.078) (Figure 3E-F), consistent with feedback reactivation of MAPK mediated by wild-type RAS isoforms. Supporting this, we observed rebound of phospho-ERK, as well as increased engagement of HRAS-GTP and NRAS-GTP with RAF over the first 48 hours of treatment with both sotorasib and BBO-8520 (Supplemental Figure 3A-C). Knock down of HRAS/NRAS expression with siRNA blunted MAPK reactivation and led to greater suppression of phospho-ERK at 48 hours (Supplemental Figure 3D-E), demonstrating that activation of wild-type HRAS/NRAS plays a significant role in MAPK reactivation after both KRAS (ON) and (OFF) inhibition in *in vitro* cell models. In contrast to phospho-ERK, AKT phosphorylation levels were consistently lower in cells treated with BBO-8520 for 14 days compared to cells treated with sotorasib (p=0.001) (Figure 3C, E, G). At shorter timepoints sotorasib and vehicle-treated cells exhibited increased phospho-AKT, but this was suppressed in BBO-8520 treated cells (Supplemental Figure 3B, F), suggesting that improved suppression of KRAS^G12C^ by BBO-8520 results in decreased PI3K activation. To test this, we stably expressed AKT1^WT^ or AKT1^E17K^ in H23 cells. The AKT1^E17K^ mutation results in constitutive localization of AKT to the plasma membrane, decoupling AKT phosphorylation from upstream RAS-PI3Kα or RTK-PI3K activity^21^. In H23-AKT1^WT^ and parental H23 cells, BBO-8520 suppressed phospho-AKT to a greater extent than sotorasib, whereas BBO-8520 had no effect on pAKT levels in H23-AKT1^E17K^ cells in which AKT is decoupled from KRAS-PI3Kα (Supplemental Figure 3G). Collectively, these results indicate that BBO-8520 achieves more durable suppression of KRAS^G12C^ compared to sotorasib, which is associated with decreased PI3K-AKT activity.

### Breaking the interaction between Ras isoforms and PI3Kα improves activity of sotorasib against KRAS mutant NSCLC

Prior studies by us and others have shown that in some *KRAS^G12C^*-mutant NSCLC and CRC models, PI3K-AKT signaling increases after treatment with first-generation KRAS^G12C^ (OFF) inhibitors, and that inhibition of PI3K or downstream mediators can increase the sensitivity of *KRAS^G12C^*-mutant cells to KRAS^G12C^ (OFF) inhibition^13,17,22^. We hypothesized that the ability of BBO-8520 to better suppress KRAS^G12C^ and downstream PI3Kα-AKT signaling might explain its more potent antiproliferative effects over time. To test this, we first examined whether inhibition of reactivated KRAS-GTP in sotorasib-treated cells could suppress PI3K-AKT activity. After sotorasib treatment for 14 days, switching to BBO-8520 modestly decreased phospho-AKT levels (Supplemental Figure 4A). To determine whether this decrease in phospho-AKT was functionally relevant, we treated cells with sotorasib for 7 days followed by drug washout, continued sotorasib, or switch to BBO-8520. After drug switch, BBO-8520 significantly suppressed proliferation compared to cells maintained in sotorasib in 2 of 3 cell lines (Figure 4A, Supplemental Figure 4B-C). To validate that decreased KRAS^G12C^-mediated PI3Kα-AKT signaling was responsible for the suppression of proliferation by BBO-8520, we treated cells with sotorasib for 7 days and then added 1 μM BBOT5, a breaker tool compound that binds to the RAS-binding domain (RBD) of PI3K p110α and prevents RAS-driven activation^20^. While disrupting the RAS-p110α interaction in the absence of KRAS^G12C^ inhibition had minimal effect on cell proliferation, addition of BBOT5 in the continued presence of sotorasib suppressed proliferation to an identical degree as switching to BBO-8520 (Figure 4A, Supplemental Figure 4B-C) and was associated with partial suppression of phospho-AKT (Supplemental Figure 4D).

**Figure 4.**
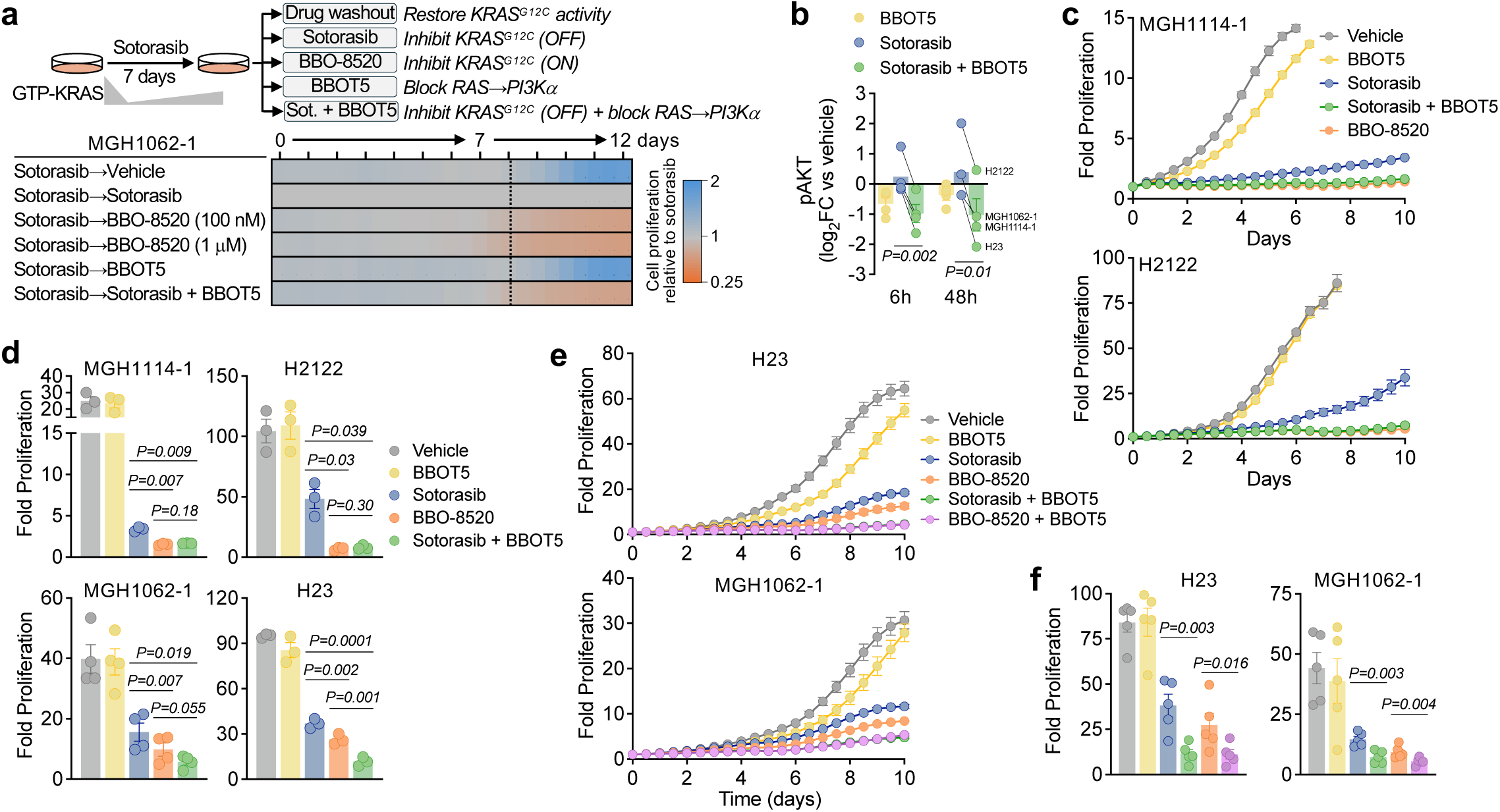
Disruption of RAS-PI3Kα increases sensitivity to sotorasib, phenocopying BBO-8520. **a,** MGH1062-1 cells were treated with sotorasib for 7 days followed by switch to the indicated conditions (sotorasib, 1 μM; BBOT5, 1 μM). Cell numbers were quantified by live cell imaging (Incucyte) and normalized to sotorasib-treated cells for each timepoint. **b,** pAKT levels after treatment with sotorasib (1 μM), BBOT5 (1 μM) or the combination were determined by western blot band quantification. Each dot represents the mean of an individual cell line, n=3 independent biological replicates. **c,** Cell were treated with sotorasib (1 μM), BBOT5 (1 μM), BBO-8520 (100 nM) or the indicated combination and cell numbers were quantified by live cell imaging (Incucyte). Data is mean and S.E.M. of n=3-4 technical replicates and is representative of n=3-4 independent biological replicates. **d,** Comparison of cell proliferation after treatment as described in panel C. Data points represent mean cell proliferation relative to baseline determined after 10 days from 3-4 independent biological replicates. **e,** Cell were treated with sotorasib (1 μM), BBOT5 (1 μM), BBO-8520 (100 nM) or combinations and cell numbers were quantified by live cell imaging (Incucyte). Data is mean and S.E.M. of n=3 technical replicates and is representative of n=5 independent biological replicates. **f,** Comparison of cell proliferation after treatment as described in panel E. Data points represent mean cell proliferation relative to baseline determined after 10 days from 5 independent biological replicates.

Next, we tested whether upfront disruption of RAS-mediated PI3Kα activation in sotorasib-treated cells might phenocopy the effects of BBO-8520 on PI3Kα signaling and proliferation. Compared to sotorasib alone, which had minimal effect or in some cases induced phospho-AKT, BBOT5 and the combination of sotorasib + BBOT5 suppressed phospho-AKT levels after 6 and 48 hours (p=0.002, 6h; p=0.01, 48h) (Figure 4B, Supplemental Figure 4E). To confirm that this decrease in phospho-AKT was due to inhibition of KRAS-mediated activation of PI3Kα, rather than blocking an interaction between wild-type HRAS/NRAS isoforms and PI3Kα, we knocked down expression of HRAS/NRAS with siRNA. phospho-AKT levels after treatment with sotorasib and sotorasib + BBOT5 were not significantly different between knockdown and non-targeting control, demonstrating that HRAS and NRAS do not significantly contribute to phospho-AKT signaling in these cell lines (Supplemental Figure 4F-G). We then examined whether the decrease in phospho-AKT signaling in sotorasib + BBOT5 treated cells was associated with decreased proliferation. While BBOT5 had minimal effect on cell proliferation alone, the combination of sotorasib + BBOT5 achieved comparable durable efficacy to BBO-8520 in 3 of 4 cell lines (Figure 4C-D, Supplemental Figure 4H), and greater suppression of proliferation in the fourth.

These results suggest that improved inhibition of KRAS^G12C^-mediated activation of PI3Kα explains the improved efficacy of BBO-8520 compared to sotorasib, however we noted that BBO-8520 or the addition of BBOT5 only partially suppressed phospho-AKT (Figure 4B, Supplemental Figure 4A,D,E), suggesting that additional mechanisms may contribute to PI3K activation. The PI3K-AKT pathway can be activated via RAS-independent activation of PI3Kα by RTKs in physiologic and oncogenic contexts^23,24^, as well as other PI3K isoforms (PI3K p110β, δ, γ)^25^. To determine whether complete suppression of phospho-AKT would further suppress cell viability, we treated cells with alpelisib (catalytic PI3K inhibitor) or BBOT5 either alone or in combination with sotorasib. In contrast to BBOT5, alpelisib fully suppressed phospho-AKT (Supplemental Figure 4I). Surprisingly, complete suppression of phospho-AKT with sotorasib + alpelisib did not translate into greater suppression of cell viability over time compared to sotorasib + BBOT5 (Supplemental Figure 4J). This suggests that although KRAS^G12C^-driven activation of PI3Kα generates only part of the total cellular pool of phosphorylated AKT, it represents the main source of oncogenic PI3Kα-AKT signaling in these models and indicates that relative changes in total AKT phosphorylation may underestimate the effect on cell proliferation/viability. Finally, given the greater suppression of cell proliferation that we observed in H23 and MGH1062-1 cell lines with the combination of sotorasib + BBOT5 compared to BBO-8520 (Figure 4D, Supplemental Figure 4H), we tested whether we could further sensitize these cell lines by combining BBOT5 with BBO-8520. Indeed, we observed greater suppression of cell proliferation when combining BBO-8520 with BBOT5 compared to BBO-8520 alone in both models (Figure 4E-F). These results suggest that in these two cell lines, disrupting the interaction between wild-type KRAS (since these cell lines are heterozygous for KRAS^G12C^) or non-canonical RAS proteins (e.g., non-HRAS, NRAS) and PI3Kα may further improve the efficacy of KRAS^G12C^ (ON) inhibitors.

Collectively, these results demonstrate that the KRAS^G12C^ (ON) inhibitor BBO-8520 achieves more durable suppression of KRAS compared to the KRAS^G12C^ (OFF) inhibitor sotorasib. Although deep and durable suppression of downstream MAPK signaling is limited by bypass reactivation by wild-type HRAS and NRAS in these cell models, improved suppression of KRAS^G12C^-driven PI3Kα-AKT, which is not reactivated by HRAS/NRAS, leads to more durable suppression of proliferation. Disruption of the KRAS^G12C^-p110α interaction can decrease KRAS-driven oncogenic PI3Kα-AKT signaling and improve the efficacy of sotorasib, and in some cases, suppression of PI3Kα activation by RAS family proteins other than HRAS/NRAS may additionally improve the efficacy of sotorasib and BBO-8520.

### Disruption of RAS-PI3Kα potentiates *in vivo* activity of sotorasib and BBO-8520

We next compared the antitumor activity of BBO-8520 and sotorasib in murine xenograft models with diverse co-occurring mutations (Supplemental Figure 1A). In the MGH9029-1 PDX model (generated from the same patient as the MGH1114-1 cell line), both BBO-8520 and sotorasib induced tumor regression, with greater tumor regression after treatment with 30 mg/kg BBO-8520 compared to 100 mg/kg sotorasib (Figure 5A-B, Supplemental Figure 5A-B), including 5/15 complete responses. In the MGH1138-1 PDX model, which is less sensitive to sotorasib^26^, 10 mg/kg BBO-8520 induced tumor regression whereas 100 mg/kg sotorasib induced partial tumor growth inhibition but not regression (Figure 5C-D, Supplemental Figure 5C-D). Both of these KRAS^G12C^ PDX models harbored co-occurring STK11 mutations which highlight BBO-8520’s potential benefit in this underserved patient population. The free drug concentration was 69- and 22-fold higher with sotorasib (free AUC_0-24h_ of 241.1 h*ng/mL with 100 mg/kg sotorasib) compared to 10 mg/kg and 30 mg/kg BBO-8520, respectively, (free AUC_0-24h_ of 3.5 and 10.8 h*ng/mL with 10 and 30 mg/kg BBO-8520, respectively) in a PK study in the same strain of mice, indicating that BBO-8520 achieves greater efficacy despite markedly lower free drug exposure (Supplemental Figure 5E). Notably, consistent with these preclinical findings, early clinical PK data also show that BBO-8520 exhibits significantly greater monotherapy activity at substantially lower free drug concentrations than KRAS (OFF) inhibitors^18^. Next, we tested whether disruption of the RAS-PI3Kα interaction could increase the response to sotorasib. Consistent with our *in vitro* observations, addition of the clinical RAS-PI3Kα breaker BBO-10203 to sotorasib induced strong tumor regressions in the MGH1138-1 and H2122 models (Figure 5C-D, Supplemental Figure 5F). Interestingly, in the MGH1138-1 model, the combination achieved very similar efficacy as BBO-8520 monotherapy. In contrast, adding BBO-10203 to a maximal efficacious dose of BBO-8520 (30 mg/kg) in the MGH1138-1 model did not lead to better efficacy (Supplemental Figure 5G-H), supporting the notion that KRAS*^G12C^* (ON) can be the sole driver of both the MAPK and PI3Kα signaling pathways.

**Figure 5.**
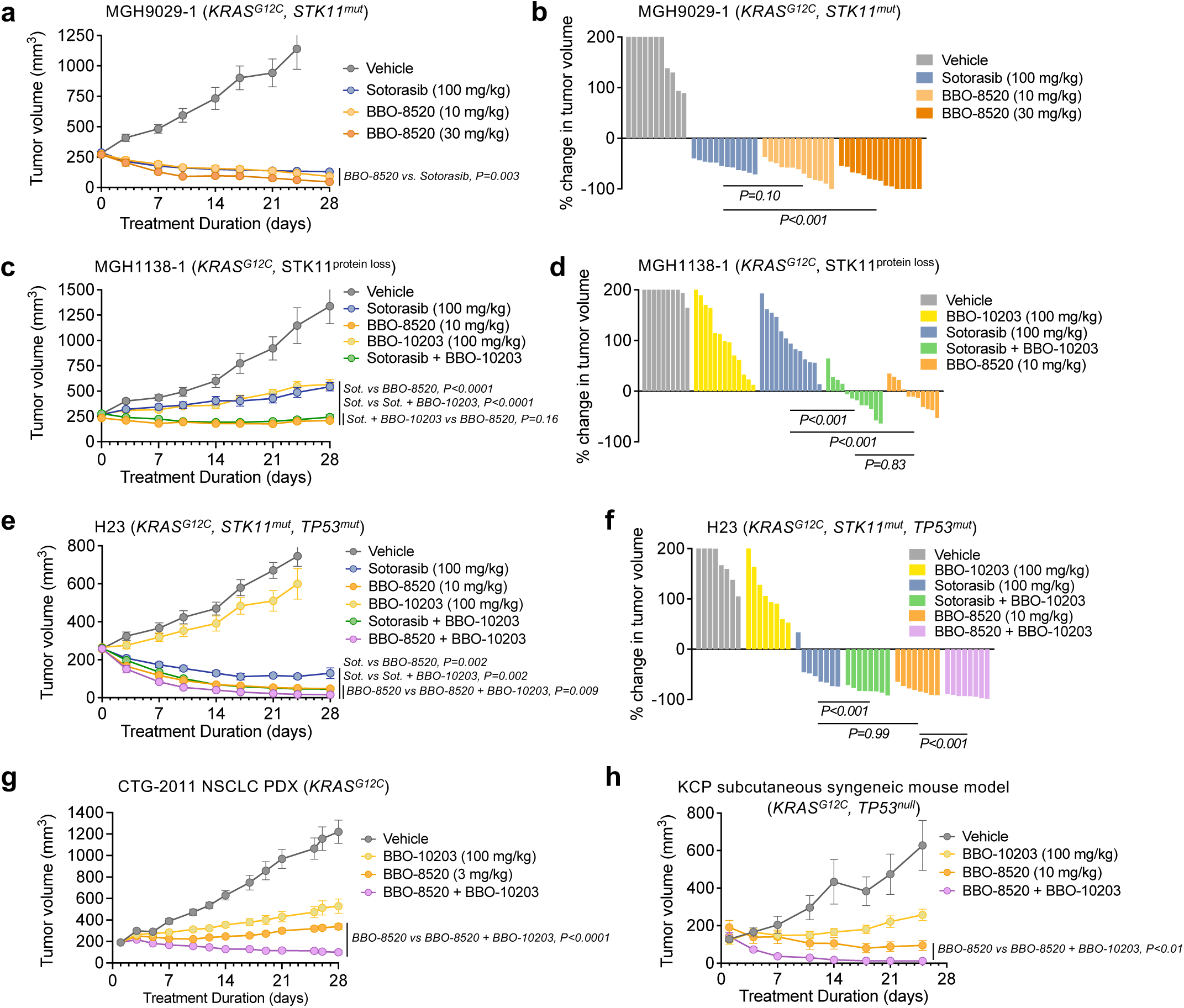
Disruption of RAS-PI3Kα with BBO-10203 increases *in vivo* efficacy of KRAS (OFF) and KRAS (ON) inhibitors. **a,** Mice bearing MGH9029-1 PDX tumors were treated with vehicle (n=11), sotorasib (100 mg/kg, n=12), BBO-8520 (10 mg/kg, n=13; 30 mg/kg, n=15) once daily by oral gavage. Data are mean and S.E.M. **b,** Waterfall plot of the change in tumor volume at 28 days compared to baseline (except vehicle for which final volumes were recorded at 24 days). **c,** Mice bearing MGH1138-1 PDX tumors were treated with vehicle (n=10), sotorasib (100 mg/kg, n=13), BBO-8520 (10 mg/kg, n=11), BBO-10203 (100 mg/kg, n=13), or sotorasib + BBO-10203 (n=12) once daily by oral gavage. Data are mean and S.E.M. **d,** Waterfall plot of the change in tumor volume at 28 days compared to baseline. **e,** Mice bearing H23 xenograft tumors were treated with vehicle (n=8), sotorasib (100 mg/kg, n=8), BBO-8520 (10 mg/kg, n=11), BBO-10203 (100 mg/kg; n=13), sotorasib + BBO-10203 (n=8) or BBO-8520 + BBO-10203 (n=8) once daily by oral gavage. Data are mean and S.E.M. **f,** Waterfall plot of the change in tumor volume at 28 days compared to baseline (except vehicle and BBO-10203 for which final volumes were recorded at 24 days). **g,** Mice bearing CTG-2011 PDX tumors were treated with vehicle (n=10), BBO-8520 (3 mg/kg, n=10), BBO-10203 (100 mg/kg; n=10) or BBO-8520 + BBO-10203 (n=10) once daily by oral gavage. Data are mean and S.E.M. **h,** Mice bearing sub-cutaneous KCP tumors were treated with vehicle (n=5), BBO-8520 (10 mg/kg, n=8), BBO-10203 (100 mg/kg; n=8) or BBO-8520 + BBO-10203 (n=8) once daily by oral gavage. Data are mean and S.E.M.

While our results demonstrate that KRAS (ON) inhibitors achieve better suppression of KRAS^G12C^ -activated PI3Kα, compared to KRAS (OFF) inhibitors, our current *in vitro* results along with our recently reported work^20^ suggest that further disruption of interactions between RAS proteins and PI3Kα might further improve the antitumor efficacy of KRAS (ON) inhibitors in some contexts. To test this, we established H23 xenograft tumors in mice. While sotorasib (100 mg/kg) induced tumor regression, the addition of BBO-10203 further deepened the response (Figure 5E-F, Supplemental Figure 5I), achieving tumor regression levels comparable to that observed with BBO-8520 monotherapy (10 mg/kg). These combination results were consistent with those observed in the MGH1138-1 model. However, adding BBO-10203 to BBO-8520 (10 mg/kg) in the H23 model did enhance the depth of tumor regression (Figure 5E-F) demonstrating the benefit of additional suppression of RAS-PI3K in combination with KRAS (ON) inhibition. We observed similar combination benefit in the CTG-2011 NSCLC PDX model (Figure 5G, Supplemental Figure 5J) as well as the subcutaneous KCP (*Kras^G12^*^C^, *Trp53^null^*) syngeneic mouse model (Figure 5H, Supplemental Figure 5K). Moreover, in the autochthonous KCP syngeneic mouse model, which is highly sensitive to BBO-8520 monotherapy, we observed that after 6 weeks of treatment followed by drug removal for 4 weeks, the combination of BBO-8520 and BBO-10203 more durably suppressed tumor regrowth than BBO-8520 alone (Supplemental Figure 5L). Collectively, these results demonstrate that disruption of RAS-PI3Kα with BBO-10203 in combination with sotorasib phenocopies the anti-tumor activity of BBO-8520 alone and can additionally improve the durability and anti-tumor efficacy of BBO-8520 in a RAS isoform agnostic manner.

## Discussion

Here we highlight the therapeutic potential of BBO-8520, a dual-state, KRAS^G12C^(ON) and (OFF) inhibitor, which exhibits increased potency and rapid target engagement compared to first-generation KRAS^G12C^ inhibitors. At present, several strategies are being pursued to improve upon the clinical activity of sotorasib and adagrasib. Next-generation (OFF)-only inhibitors (divarasib, D3S-001) exhibit improvements in potency and target engagement and have shown improved clinical efficacy, demonstrating the clinical potential of more effective targeting of KRAS-GDP^27,28^. On the other hand, targeting the GTP-bound KRAS^G12C^ (ON) state enables inhibition of signaling competent KRAS and does not rely on intrinsic GTP hydrolysis. In contrast to cyclophilin A-binding tri-complex inhibitors that selectively bind to the GTP-bound (ON) state, BBO-8520 irreversibly binds to both the (ON) and (OFF) state of KRAS^G12C^. We show that BBO-8520 exhibits, in addition to superior potency, a greater magnitude and durability of suppression of signaling-competent KRAS^G12C^ compared to the KRAS^G12C^ (OFF)-only inhibitor, sotorasib.

The efficacy of first-generation KRAS^G12C^ inhibitors is limited by adaptive feedback reactivation of the MAPK pathway that can result from RTK-driven activation of wild-type RAS isoforms^14,17^ as well as via newly synthesized KRAS^G12C^ that adopts a GTP-bound state^12^. Whether one or both of these mechanisms dominates, and whether this is context-specific, remains an open question. In principle, targeting the GTP-bound KRAS^G12C^ (ON) state would be expected to overcome the latter, but still be susceptible to feedback reactivation driven by the former. In the lung cancer cell lines examined in this study, despite more durable suppression of signaling-competent KRAS^G12C^ (e.g. GTP-bound KRAS^G12C^ capable of binding to RBD-RAF), we observed similar feedback reactivation of MAPK regardless of treatment with sotorasib or BBO-8520. Moreover, in both instances, this was reversed by knock-down of HRAS/NRAS, demonstrating a dominant role for wild-type RAS isoforms in driving adaptive MAPK reactivation in this *in vitro* context, which cannot be overcome by targeting KRAS^G12C^ (ON). The role of WT RAS isoform reactivation *in vivo* remains to be understood, as KRAS^G12C^ inhibition can lead to sustained monotherapy activity in tumor models. In contrast, we observed that BBO-8520 decreased PI3Kα-AKT signaling compared with sotorasib, and HRAS/NRAS knock-down did not reduce PI3Kα-AKT activation in sotorasib-treated cells. These results indicate that although oncogenic KRAS^G12C^ and wild-type RAS isoforms are all effective in activating MAPK signaling, oncogenic KRAS appears uniquely capable of activating PI3Kα. As a result, wild-type HRAS/NRAS isoforms cannot fully compensate for the loss of oncogenic KRAS, such that deeper and more durable suppression of KRAS^G12C^ (ON) by BBO-8520 leads to reduced PI3Kα-AKT activation. Collectively, these results demonstrate a role for feedback reactivation of both KRAS^G12C^ and wild-type RAS isoforms and reveal distinct signaling outcomes that result from respective differences in their ability to activate effector proteins.

Our study adds to prior work demonstrating the therapeutic importance of suppression of the PI3Kα-AKT pathway in *KRAS*-mutant cancer cells. PI3Kα and AKT inhibitors have been shown to improve the efficacy of MEK and first-generation KRAS^G12C^ inhibitors^13,29–31^. PI3Kα can be activated by RAS-independent and RAS-dependent mechanisms in a context specific manner. In many normal physiologic systems, canonical PI3Kα-AKT signaling is activated via receptor tyrosine kinases (e.g. in response to insulin or growth factors) independent of RAS^32^. On the other hand, particularly in the context of cancer, RAS has been shown to directly bind and activate PI3K p110α, driving transformation and tumorigenesis^33–35^. Within cancer, the relative contribution of RAS-dependent and independent regulation of PI3Kα may be context dependent. In *KRAS* and *EGFR*-driven lung cancer mouse models, genetic disruption of the RAS-binding domain of p110α impaired PI3K signaling and inhibited tumor growth^35,36^. On the other hand, other studies have suggested a more limited role for direct activation PI3Kα by RAS in contexts such as colon cancer^37^ and pancreatic cancer^38^. Our results suggest that better suppression of PI3Kα-AKT downstream of KRAS^G12C^ signaling by BBO-8520 compared to sotorasib, or by sotorasib + BBO-10203 compared to sotorasib alone, explains the improved suppression of proliferation over time. This is consistent with a recent study demonstrating that sensitivity of *KRAS^G12C^*-mutant cell lines to KRAS^G12C^ inhibitors correlates with the degree of inhibition of PI3Kα-AKT signaling^22^. Interestingly, we note that the decrease in pAKT after treatment with BBO-8520 or BBOT5 is partial and relatively modest. By comparison, the catalytic PI3Kα inhibitor alpelisib completely suppressed pAKT, suggesting that in these cells, total pAKT levels reflect both RAS-dependent and independent PI3Kα activation. Despite this difference in pAKT suppression, we observed nearly identical effects on cell viability when combining BBOT5 or alpelisib with sotorasib, suggesting that the overall cellular pAKT level may reflect functionally distinct PI3Kα-AKT signals: 1) oncogenic RAS-dependent PI3Kα activation that drives cell proliferation and survival and 2) RAS-independent PI3Kα-AKT signaling that is largely dispensable for the tumor cell but drives dose limiting hyperglycemia in patients

We consistently observed that combining BBOT5/BBO-10203 with sotorasib recapitulated the activity of BBO-8520, achieving greater inhibition of cell proliferation and tumor regression, thus demonstrating the importance of fully suppressing RAS-PI3Kα signaling in *KRAS^G12C^*-mutant NSCLC. Interestingly, in a subset of models (MGH1114-1 and MGH1138-1) adding BBO-10203 to BBO-8520 at maximal dose - 30 mg/kg) did not enhance efficacy suggesting that in these models, KRAS^G12C^ (ON) is the dominant regulator of oncogenic RAS-PI3Kα signaling. In other models, however, (H23, MGH1062-1, CTG-2011, KCP) co-treatment with BBOT5/BBO-10203 + BBO-8520 or sotorasib led to greater antitumor activity than BBO-8520 alone, suggesting that additional RAS-input into PI3Kα augments KRAS^G12C^, extending recently reported results in the H2122, H358 and SW1573 NSCLC models demonstrating the benefit of combining BBO-10203 with BBO-8520 and other oncogene-targeted therapies^20^. We did not observe any impact of HRAS/NRAS knockdown on PI3Kα-AKT signaling in this context, suggesting that disruption of interactions between PI3Kα and non-canonical RAS proteins or wild-type KRAS (in heterozygous tumor contexts) may contribute to efficacy of BBOT5/BBO-10203 in some cases. Further studies will be required to elucidate the underlying mechanisms that contribute to heterogeneity of RAS-dependent PI3Kα activation in NSCLC. For example, wild-type HRAS/NRAS driven PI3Kα signaling may contribute in some contexts, as recently demonstrated in *KRAS^G12C^*-mutant cells^39^. BBO-10203 and other recently described RAS-p110a inhibitors (e.g., VVD-159642)^40^ will be valuable tools for further understanding the role of RAS-p110α interactions in cancer.

Overall, the work presented here offers insights into the regulation of PI3Kα signaling by oncogenic and wild-type RAS isoforms and highlights the importance of suppressing RAS-PI3Kα-AKT in *KRAS*-mutant lung cancer. We demonstrate that BBO-8520 achieves more durable suppression of signaling-competent GTP-bound KRAS^G12C^ and downstream PI3Kα-AKT signaling, offering a mechanistic explanation for the improved efficacy of dual-state KRAS^G12C^ inhibition. Further, our work lends further support for the potential for BBO-10203 and other breakers of the RAS-PI3Kα interaction in the treatment of cancers dependent on oncogenic RAS-PI3Kα signaling in combination with KRAS inhibitors.

## Materials and Methods

### Cell lines and cell culture

All cell lines are listed in Supplementary Figure 1A. H23, H358, H1792, H2030, H2122, HCC44, LU65, and LU99A cells were obtained from the MGH Center for Molecular Therapeutics. The identities of these cell lines were verified by STR analysis (Bio-synthesis, Inc.) at the time that these studies were performed. Patient-derived cell lines (MGH1062-1, MGH1088-1, MGH1112-1, MGH1114-1, MGH1138-1, and MGH1143-2) were established in our laboratory from core biopsy or pleural effusion samples as previously described^26,41^. All patients signed informed consent to participate in a Dana-Farber–Harvard Cancer Center Institutional Review Board–approved protocol giving permission for research to be performed on their samples. All cell lines were maintained in RPMI (Life Technologies) supplemented with 10% FBS (Life Technologies).

### Inhibitors

Sotorasib (MedChemExpress) was diluted in DMSO to a concentration of 10 mM and stored at - 20°C until immediately prior to use. BBO-7410, BBO-8520, BBOT5, BBO-10203 were provided by BridgeBio Oncology Therapeutics and were diluted in DMSO to 10 mM and stored at −20°C until immediately prior to use.

### Cell proliferation assays

For short-term assays, cells were plated in 96-well plates at a concentration of 3,000 cells per well. Cells were allowed to adhere overnight and were then treated with a 9-point titration starting at 0.01 nM (BBO-8520, BBO-7410) or 0.1 nM (sotorasib) and increasing at half-log intervals using a Tecan D300e Digital Dispenser. Cell proliferation was determined by CellTiter-Glo (Promega) after 72 hours using a 1:4 reagent:media ratio; luminescence was measured at 560 nm on a Spectramax I3x spectrophotometer (Molecular Devices). IC_50_ and E_max_ were calculated using GraphPad Prism Software using the function log(inhibitor) versus response-variable slope (4 parameters). For long-term assays, cells were transduced with lentivirus containing Nuclight green (Essen Biosciences) and selected in puromycin. Cells were seeded in 96-well plates at a concentration of 300 to 1,000 cells per well. Cells were allowed to adhere overnight and were treated the following day with inhibitors. Cell counts were performed at 6- or 12-hour intervals using IncuCyte live cell imaging. Fresh media and inhibitors were exchanged twice per week. Data was plotted using GraphPad Prism software. For fold proliferation values, each timepoint cell count was normalized to the cell count at day 0.

### Viral transduction

For Nuclight green (Essen Biosciences) viral transduction, cells were seeded in 12-well plates and allowed to adhere overnight. On the following day, 10.4 μg/mL virus and 8 μg/mL polybrene were added to fresh growth medium. Virus + polybrene containing media were added to cells, and plates were spun at 1200rpm, 32C for 1 hour. Cells were selected with 2ug/mL puromycin.

For the AKT1^wt^ and AKT1^E17K^ lentiviral production, all plasmid inserts were provided by Dr. Alex Toker of Harvard Medical School. HEK293FT cells were plated in 10 cm plates and allowed to adhere overnight in DMEM supplemented with 10% FBS. Cells were transfected according to the manufacturer’s recommendation using 3 mL of OptiMEM combined with 41 μL Lipofectamine 3000, 35 μL P3000, 8.7 μg psPAX2, 4.3 μg pMD2G, and 4.3 μg of the target construct. Cells were incubated for 6 hours at 37°C prior to changing media to DMEM + 10% FBS + pen/strep (complete D10). 24 hours later, 12 mL of supernatant was collected and stored at 4°C, and 12 mL of fresh, complete D10 was added to the plate. The final 12 mL of media was collected 24 hours later and pooled with the first harvest. 6 mL of 5X PEG-it was added to 24 mL of supernatant and allowed to incubate for 24 hours at 4°C. The following day the solution was centrifuged at 1500 x g for 30 minutes at 4°C. The virus was resuspended in 300 μL PBS and stored at −80°C.

For viral transduction, H23 cells were seeded in 6-well plates and allowed to adhere overnight. The following day, 8 μg/mL of polybrene was added to the wells. Virus was then added at a range of volumes, and plates were spun at 1500 x g for 30 minutes at 4°C. After 6 hours incubation at 37°C, 2mL of fresh media was added to each well. Cells were selected the following day for 5 days using 500 μg/mL G418 until all non-transduced cells were dead.

### Western blotting and antibodies

Cells were seeded in 6-well or 10 cm plates and allowed to adhere overnight prior to treatment. For short-term experiments, cells were treated the following day with the indicated inhibitors for 6, 24, or 48 hours. For long-term experiments, cells were cultured in the indicated inhibitor for 2 weeks prior to seeding. Cells were maintained at 80% confluence or below and fresh media and inhibitors were added twice per week. After treatment, cells were washed with ice cold PBS and lysates were collected with lysis buffer (20 mM TRIS-HCl pH 7.4, 150 mM NaCl, 1% NP-40 Surfact-Amps, 1 mM EDTA pH 8.0, 1 mM EGTA, 10% glycerol) supplemented with 5 mM NaPPi, 50 mM NaF, 10 mM beta-glycerol phosphate, 2 mM NaVO3, 1 mM PMSF, 16 μg/ml aprotinin, 16 μg/ml leupeptin, 16 μg/ml pepstatin, and 0.5mM DTT. Lysates were centrifuged to remove cell debris and protein levels in supernatants were quantified via BCA assay (Thermo Scientific). Protein electrophoresis was performed using 4-12% NuPAGE Bis-Tris gels (Thermo Scientific) in MOPS SDS running buffer (Thermo Scientific) before transferring to PVDF membranes (ThermoFisher). Membranes were blocked in 5% milk/TBS-T and then incubated with primary antibody. All primary antibodies were diluted in 5% bovine serum albumin as follows: KRAS 1:2000 (Sigma WH0003845M1), HRAS 1:500 (Proteintech 18295-1-AP), NRAS 1:500 (Santa Cruz Biosciences sc-31), phospho-p44/42 MAPK (Erk1/2) Thr202/Tyr204 1:1,000 (Cell Signaling Technology CST9101), p44/42 MAPK (Erk1/2) 1:1,000 (Cell Signaling Technology CST9102), phospho-RSK1 T359+S363, 1:1,000 (Abcam ab32413), phospho-AKT Ser473, 1:1,000 (Cell Signaling Technology CST4060), AKT 1:1,000 (Cell Signaling Technology CST4691), and B-actin 1:1,000 (Cell Signaling Technology CST4970). Membranes were washed with TBS-T, incubated with the secondary antibody diluted 1:12500 in 5% milk at room temperature (anti-rabbit IgG Cell Signaling Technology CST7074; anti-mouse IgG Cell Signaling Technology CST7076). Membranes were washed in TBS-T and then developed using SuperSignal West Femto (Thermo Fisher). Luminescence was imaged using a G:Box Chemi-XRQ system (Syngene). Western blots were quantified using the GeneTools software and the “Manual Band Quantification” tool (automatic background correction).

### RAS-GTP Pulldown

RAS activity was assessed by GST-RAF-RBD pulldown (Cell Signaling Technology, Active Ras Detection Kit). All reagents were prepared and supplemented according to manufacturer’s instructions. Experiment was conducted according to manufacturer’s instructions. 500 μg of protein lysate was incubated with GST-RAF-RBD overnight at 4°C. After completion of the pulldown, western blot analysis was conducted according to the methods above. The pan RAS antibody was provided in the kit, and the RAS isoform–specific antibodies used were the same as listed above (KRAS 1:2000, Sigma WH0003845M1; HRAS 1:500, Proteintech 18295-1-AP; NRAS 1:500, Santa Cruz Biosciences sc-31).

### siRNA-mediated gene knockdown

siRNA transfections were performed with Lipofectamine RNAiMAX Transfection Reagent according to the manufacturer’s protocol (Invitrogen, Cat#13778075). Cells were seeded in 10 cm plates and allowed to adhere overnight. Prior to transfection, cell culture media was replaced with antibiotic-free media. 24 hours after transfection, cells were trypsinized and plated for treatment and subsequent Western blot. The following Dharmacon siRNA were used at 10nM concentration: Non-targeting control (NTC) (D-001810-01-05), NRAS (L-003919-00-0005), and HRAS (L-004142-00-0005).

### Mouse studies

All *in vivo* procedures were reviewed and approved by the Institutional Animal Care and Use Committee (IACUC) prior to execution and performed in accordance with the regulations and guidelines of the Association for Assessment and Accreditation of Laboratory Animal Care at MGH, NYU, Champions Oncology, Inc., or Crown Biosciences, Inc.

*KRAS*-mutant NSCLC PDX models were generated by subcutaneous implantation of tumor cells or tissue from malignant pleural effusions, core-needle biopsies, or surgical resections into male NSG mice (Jackson Labs). For models generated at MGH, all patients signed informed consent to participate in Dana Farber/Harvard Cancer Center IRB-approved protocol giving permission for research to be conducted on their sample. All animal studies were conducted through MGH IACUC-approved animal protocols in accordance with institutional guidelines. Subcutaneous tumors were serially passaged twice to fully establish each model. All mice used were between 8 weeks and 6 months of age. Mice were kept at a temperature of 74 ± 2 °F (23 ± 1 °C) with a 12-h light/12-h dark cycle. Relative humidity was maintained at 30–70%. Maximum tumor size did not exceed 2000 mm^3^, in accordance with IACUC regulations. For drug treatment studies, PDX tumor tissue was directly implanted (MGH9029-1, MGH1138-1, CTG-2011) or cells were injected as a 1:1 matrigel:PBS suspension (H23, H2122) unilaterally into the subcutaneous space on the flank of immunocompromised mice or the murine SPC-Cre KrasG12C/+;p53fl/fl (KCP) cell line was injected unilaterally into the subcutaneous space on the flank of immunocompetent mice. Tumors were measured with electronic calipers, and the tumor volume was calculated according to the formula V = 0.52 × L × W^2^. Mice with established tumors (150–350 mm^3^) were randomized to drug treatment groups to minimize differences in baseline tumor volumes. For the NSCLC autochthonous GEMM efficacy study, KRAS^G12C^;Tp53^R270H^ (KCP) mice^42^ (mixed background) were monitored by MRI for tumor development after intranasal induction with adeno-Cre (2.5 × 10^6^ PFU). When lung tumors reached a mean size of 84 mm^3^, mice were randomized into treatment groups (n=10 per group) and orally dosed daily with vehicle or 10 mg/kg BBO-8520 for 6 weeks. Lung tumor volume was monitored by MRI every 2 weeks for 10 weeks. BBO-8520 and sotorasib were formulated in 10% v/v N-methyl-pyrrolidone (Sigma-Aldrich, catalog 328634), 20% w/v solutol (Sigma-Aldrich, catalog 8074841000), and 30% v/v polyethylene glycol 300 (Sigma-Aldrich, catalog 202371) in 50-mmol/L citrate buffer pH 4 to 5. BBO-10203 was formulated in 0.5% v/v Tween 80 (Sigma-Aldrich, catalog A43489), 0.5%w/v methylcellulose (Sigma-Aldrich, catalog M0262-1KG) dissolved in purified sterile water). Vehicle (formulation buffer) was administered once daily by oral gavage, 7 days per week. BBO-8520 was administered once daily by oral gavage at 10 mg/kg or 30 mg/kg once daily, 7 days per week. Sotorasib and BBO-10203 were administered once daily by oral gavage at 100 mg/kg, 7 days per week.

For the NSG mouse PK study, blood was collected from 3 animals in each group 2, 6, and 24 hours following the final a single oral dose of 10 mg/kg BBO-8520, 30 mg/kg BBO-8520, or 100 mg/kg sotorasib, which were formulated as described above. Blood was collected via the tail vein into BD Microcontainer tubes with K2 EDTA (BD, catalog 367841) and the plasma was collected according to manufacturer’s instructions and processed into plasma by centrifuging at 4°C for 5,000 x g for 10 minutes and then stored at −80°C. Protein precipitated plasma samples were analyzed with liquid chromatography tandem mass spectrometry at Cytoscient, LLC using a Sciex 5500 and analyzed with Analyst Software 1.7.1. Total AUC_(0-24h)_ were calculated from linear trapezoidal linear interpolation integration of plasma concentrations generated from composite sampling of mice. Free fraction adjusted AUC_(0-24h)_ were calculated by multiplying total AUC_(0-24h)_ by the fraction unbound (fu) in mouse plasma for each compound (BBO-8520 fu = 0.0026; sotorasib fu = 0.06).

### Statistical analysis

Data analyses were performed using Microsoft Excel or GraphPad Prism software (version 10). Two tailed paired t-tests were used when 1) comparing the differences between the IC_50_ and E_max_ of sotorasib and BBO-8520 in individual models, 2) comparing the effect of BBO-8520, sotorasib, and combination treatments on long-term viability, and 3) comparing the effect of BBO-8520, sotorasib, and combination treatments on downstream signaling in individual models. When comparing downstream signaling levels of the models as a cohort, we used a two-tailed ratio paired t-test. When comparing the IC_50_ and E_max_ of the different compounds across all models, we used a Wilcoxon signed rank test, and we used a Mann-Whitney test when comparing IC_50_ and E_max_ of STK11 or TP53 mutant versus wild-type. For the *in vivo* efficacy combination study statistical analyses, a one-way repeated measures ANOVA was first performed to confirm that there was a statistically significant difference between the group mean tumor volumes and then statistical analyses were performed to compare the monotherapy and combination group means. Two-way repeated-measures ANOVA was performed between the indicated group means over the number of indicated days. A p value of less than 0.05 was considered statistically significant.

## Acknowledgements

This work was supported by a research grant provided by BBOT (to A.N.H and to K.K.W.) and by the Ludwig Center at Harvard. This project has been funded in part with Federal funds from the National Cancer Institute, National Institutes of Health, under Contract No. 75N91019D00024. The content of this publication does not necessarily reflect the views or policies of the Department of Health and Human Services nor does mention of trade names, commercial products, or organizations imply endorsement by the US government. We thank A. Toker (Harvard Medical School) for the AKT1^WT^ and AKT1^E17K^ expression vectors. We thank members of the Hata Lab and MGH Thoracic Oncology group for helpful feedback and support.

## Supplemental Figure Legends

**Supplemental Figure 1 The KRAS^G12C^ (ON) inhibitor BBO-8520 exhibits more potent suppression of MAPK signaling and cell proliferation compared to the KRAS^G12C^ (OFF) inhibitor sotorasib. A,** *KRAS^G12C^*-mutant NSCLC cell lines used in this study. **B,** Composite dose response curves of *KRAS^G12C^*-mutant NSCLC cell lines treated with KRAS inhibitors for 72 hours. Cell viability was quantified by CellTiter-Glo. Curves shown are mean and S.E.M. of n=3-7 independent biological replicates. **C,** Comparison of IC50 values (3-day viability assays) for BBO-8520 versus sotorasib or adagrasib. Each value is the mean IC50 of n=3-7 independent biological replicates. **D,** Western blot analysis of *KRAS^G12C^*-mutant NSCLC cell lines treated with increasing concentrations of sotorasib or BBO-8520 for 6 hours. Data is representative of n=3 independent biological replicates. **E,** Comparison of IC50 and Emax values from 3-day viability assays, stratified by co-occurring mutations in TP53 or STK11. Data points are mean of n=3-7 biological replicates.

**Supplemental Figure 2. BBO-8520 exhibits more durable suppression of cell proliferation compared to sotorasib. A,** Generation of GFP-labeled cell lines for live cell imaging. Representative images of the H23 cell line before and after labeling are shown. B, Composite dose response curves of matched parental and GFP-labeled cell lines treated with KRAS inhibitors for 72 hours. Cell viability was quantified by CellTiter-Glo. GFP-labelled curves shown are mean of n=2 independent biological replicates. **C,** Imaged-based monitoring (Incucyte) of proliferation in cell lines treated with sotorasib or BBO-8520. Data are mean and S.E.M of n=4 technical replicates and are representative of n=3-4 independent biological replicates. **D,** Relative cell proliferation of cells treated with BBO-8520, normalized to sotorasib. Values are mean of n=3-4 biological replicates, corresponding to the cell counts shown in panel C. **E,** Comparison of cell proliferation after treatment with 10 nM, 100 nM or 1 μM BBO-8520 or 1 μM sotorasib. Values represent cell proliferation relative to baseline determined after 10 days (H23, MGH1062-1, H2030), 6 days (H1792) or 5 days (HCC44), from 3-4 independent biological replicates.

**Supplemental Figure 3. BBO-8520 achieves more durable suppression of KRAS^G12C^-RAF engagement and inhibition of PI3K-AKT compared to sotorasib. A,** *KRAS^G12C^*-mutant NSCLC cells were treated with BBO-8520 (100 nM) or sotorasib (1 μM) for 6 or 48 hours and RAF-RBD pull-down was performed to assess engagement between KRAS-GTP, NRAS-GTP, HRAS-GTP and RAF. Data are representative of n=3 independent biological replicates and correspond to quantified values in Figure 3A. **B,** Representative western blot images of cells treated with BBO-8520 (100 nM) or sotorasib (1 μM) for 6, 24, or 48 hours. Data are representative of n=7-8 independent biological replicates and correspond to the quantified values in panels C and E. **C,** Average change in phospho-ERK levels after treatment with sotorasib (1 μM) or BBO-8520 (100 nM) for 6 or 48 hours. Data are quantified band intensities from western blots (see panel B) normalized to vehicle control, n=7-8 independent biological replicates Each data point represents the average change in phosphor-ERK in an individual cell line. **D-E,** Cell lines with siRNA knockdown of HRAS/NRAS (KD) or non-targeting control (NTC) were treated with sotorasib (1 μM) or BBO-8520 (100nM) for 48 hours and harvested for western blotting. Data in panel D are quantified band intensities normalized to vehicle treated cells, n=3 independent biological replicates. Panel G shows representative western blot of n=3 biological replicates. **F,** Average change in phospho-AKT (S473) levels after treatment with sotorasib (1 μM) or BBO-8520 (100 nM) for 6 or 48 hours. Data are quantified band intensities from western blots (see panel B) normalized to vehicle control, n=7-8 independent biological replicates Each data point represents the average change in phosphor-AKT in an individual cell line. **G,** Western blot images of H23-AKT1^wt^ and H23-AKT1^E17K^ cells treated with BBO-8520 (100 nM) or sotorasib (1 μM) for 6 hours. Quantified band intensities of phospho-AKT (normalized to vehicle controls) are listed above the corresponding band.

**Supplemental Figure 4. Disruption of RAS-PI3Kα increases sensitivity to sotorasib, phenocopying BBO-8520. A,** Cells were treated with sotorasib (1 μM) for 14 days, followed by switch to BBO-8520 (100 nM) or continued sotorasib for 24 hours, then harvested for western blot analysis. **B,** Cell lines were treated with sotorasib (1 μM) for 7 days followed by switch to the indicated conditions (BBO-8520, 100 nM; BBOT5, 1 μM; sotorasib, 1 μM). Timepoints shown are relative to switch (“day 0”). Data are mean and S.E.M. of n=3-6 technical replicates and are representative of n=4 independent biological replicates. **C,** Comparison of cell proliferation after treatment as described in panel B. Data points represent mean cell proliferation 5 (MGH1062-1, H2122) or 13 (MGH1114-1) days after drug switch, normalized to the cell count at the time of drug switch, n=4 independent biological replicates. **D,** Cells were treated with sotorasib (1 μM) for 14 days, followed by addition of BBOT5 (1 μM) or continued sotorasib for 24 hours, then harvested for western blot analysis. **E,** Cells were treated with BBOT5 (1 μM), sotorasib (1 μM), or combination for 6, 24, or 48 hours then harvested for western blot analysis. Data are representative of n=3 biological replicates and correspond to quantified values in Figure 4B. **F-G,** Cell lines with siRNA knockdown of HRAS/NRAS (KD) or non-targeting control (NTC) were treated with BBOT5 (1 μM), sotorasib (1 μM) or the combination for 48 hours and harvested for western blotting. Data in panel F are quantified band intensities normalized to vehicle treated cells, n=3 independent biological replicates. Panel G shows representative western blot of n=3 biological replicates. **H,** Cells were treated with sotorasib (1 μM), BBOT5 (1 μM), BBO-8520 (100 nM) or the indicated combination and cell numbers were quantified by live cell imaging (Incucyte). Data is mean and S.E.M. of n=3-4 technical replicates and is representative of n=3-4 independent biological replicates. **I,** Cell lines were treated with BBOT5 (1 μM), alpelisib (1 μM), sotorasib (1 μM) or combinations for 24 hours and harvested for western blotting. Data are representative of n=3-4 biological replicates. **J,** Cells were treated with sotorasib (1 μM), BBOT5 (1 μM), alpelisib (1 μM) or the indicated combinations and cell numbers were quantified by live cell imaging (Incucyte). Data is mean and S.E.M. of n=3 technical replicates and is representative of n=3 independent biological replicates.

**Supplemental Figure 5. Disruption of RAS-PI3Ka with BBO-10203 increases in vivo efficacy of KRAS (OFF) and KRAS (ON) inhibitors. A,** Representative H&E images of MGH9029-1 xenograft tumors after 3 days treatment. **B,** Change in body weight of mice bearing MGH9029-1 xenograft tumors during drug treatment. Data are mean and S.E.M. (vehicle, n=11; sotorasib100 mg/kg, n=12; BBO-8520 10 mg/kg, n=13; BBO-8520 30 mg/kg, n=15). **C,** Representative H&E images of MGH9029-1 xenograft tumors after 3 days treatment. **D,** Change in body weight of mice bearing MGH9029-1 xenograft tumors during drug treatment. Data are mean and S.E.M. (vehicle, n=10; sotorasib 100 mg/kg, n=13; BBO-8520 30 mg/kg, n=11; BBO-10203 100 mg/kg, n=13; sotorasib + BBO-10203, n=12). **E,** Pharmacokinetic parameters of BBO-8520 and sotorasib in NSG mice. **F,** Mice bearing MGH1138-1 PDX tumors were treated with vehicle (n=10), sotorasib (100 mg/kg, n=13), BBO-8520 (30 mg/kg, n=11), BBO-10203 (100 mg/kg; n=13), or sotorasib + BBO-10203 (n=12) once daily by oral gavage. BBO-8520 30 mg/kg and BBO8520 30 mg/kg + BBO=10203 are shown; the other treatment arms are replotted from Figure 5C for comparison purposes. Data are mean and S.E.M. **G,** Waterfall plot of the change in tumor volume at 28 days compared to baseline. BBO-8520 30 mg/kg and BBO8520 30 mg/kg + BBO=10203 are shown; the other treatment arms are replotted from Figure 5D for comparison purposes. **H,** Mice bearing H2122 xenograft tumors were treated with vehicle (n=10), sotorasib (100 mg/kg, n=10), BBO-10203 (100 mg/kg; n=10), or sotorasib + BBO-10203 (n=10) once daily by oral gavage. Data are mean and S.E.M. **I,** Change in body weight of mice bearing H23 xenograft tumors during drug treatment. Data are mean and S.E.M. (vehicle, n=8; sotorasib 100mg/kg, n=8; BBO-8520 10 mg/kg, n=11; BBO-10203 100 mg/kg, n=13; sotorasib + BBO-10203, n=8; BBO-8520 + BBO-10203, n=8. **J,** Waterfall plot of the change in tumor volume of CTG-2011 xenograft tumors at 28 days compared to baseline. **K,** Waterfall plot of the change in tumor volume of KCP tumors at 28 days compared to baseline. **L,** Mice bearing autochthonous KCP tumors were treated with vehicle (n=14), BBO-8520 (10 mg/kg, n=15), BBO-10203 (100 mg/kg, n=11), or BBO-8520 + BBO-10203 (n=13) once daily by oral gavage for 6 weeks followed by 4 weeks with no treatment. Data are mean and S.E.M.

