## Supplemental Figures 1-5 for "Dual inhibition of GTP-bound (ON) and GDP-bound (OFF) KRAS^G12C^ suppresses PI3Kα and leads to potent tumor inhibition"

**a**

| Cell line | KRAS^G12C^ zygosity | Co-occurring mutations |
|---|---|---|
| H23 | Heterozygous | TP53, STK11 |
| H358 | Heterozygous | TP53 (deletion) |
| H1792 | Heterozygous | TP53 |
| H2030 | Homozygous | TP53, STK11 |
| H2122 | Homozygous | TP53, STK11 |
| HCC44 | Homozygous | TP53 |
| LU65 | Heterozygous | TP53 |
| LU99A | Heterozygous | TP53 |
| MGH1062 | Heterozygous | TP53 |
| MGH1088-1 | Heterozygous | --- |
| MGH1112-1 | Heterozygous | STK11 |
| MGH1114-1 | Heterozygous | STK11 |
| MGH1138-1 | Heterozygous | STK11 (protein loss) |
| MGH1143-2 | Heterozygous | TP53 |

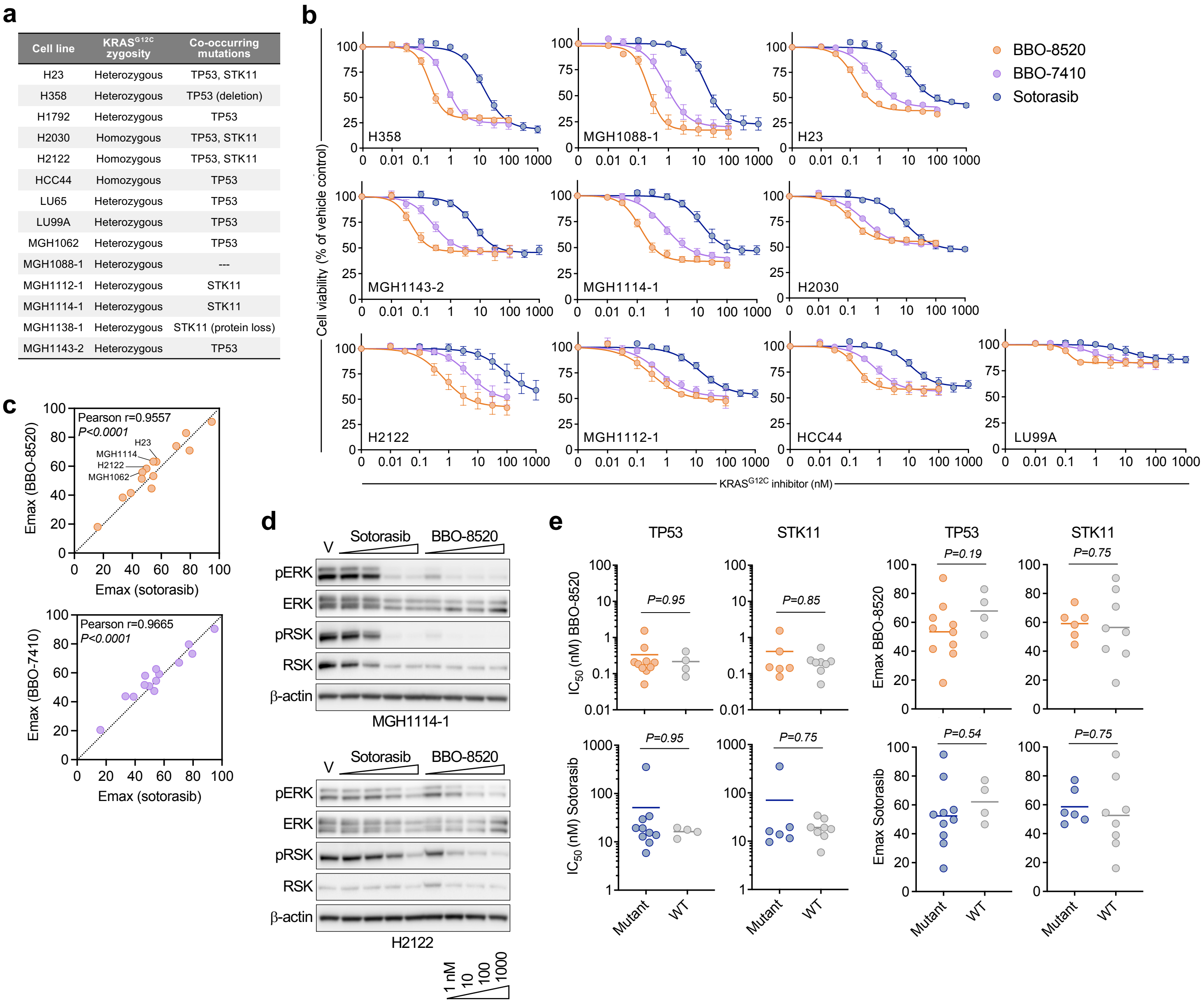


**Supplemental Figure 1. The KRAS$^{G12C}$ (ON) inhibitor BBO-8520 exhibits more potent suppression of MAPK signaling and cell proliferation compared to the KRAS$^{G12C}$ (OFF) inhibitor sotorasib. a,** *KRAS$^{G12C}$*-mutant NSCLC cell lines used in this study. **b,** Composite dose response curves of *KRAS$^{G12C}$*-mutant NSCLC cell lines treated with KRAS inhibitors for 72 hours. Cell viability was quantified by CellTiter-Glo. Curves shown are mean and S.E.M. of n=3-7 independent biological replicates. **c,** Comparison of IC50 values (3-day viability assays) for BBO-8520 or BBO-7410 versus sotorasib. Each value is the mean IC50 of n=3-7 independent biological replicates. **d,** Western blot analysis of *KRAS$^{G12C}$*-mutant NSCLC cell lines treated with increasing concentrations of sotorasib or BBO-8520 for 6 hours. Data is representative of n=3 independent biological replicates. **e,** Comparison of IC50 and Emax values from 3-day viability assays, stratified by co-occurring mutations in TP53 or STK11. Data points are mean of n=3-7 biological replicates.

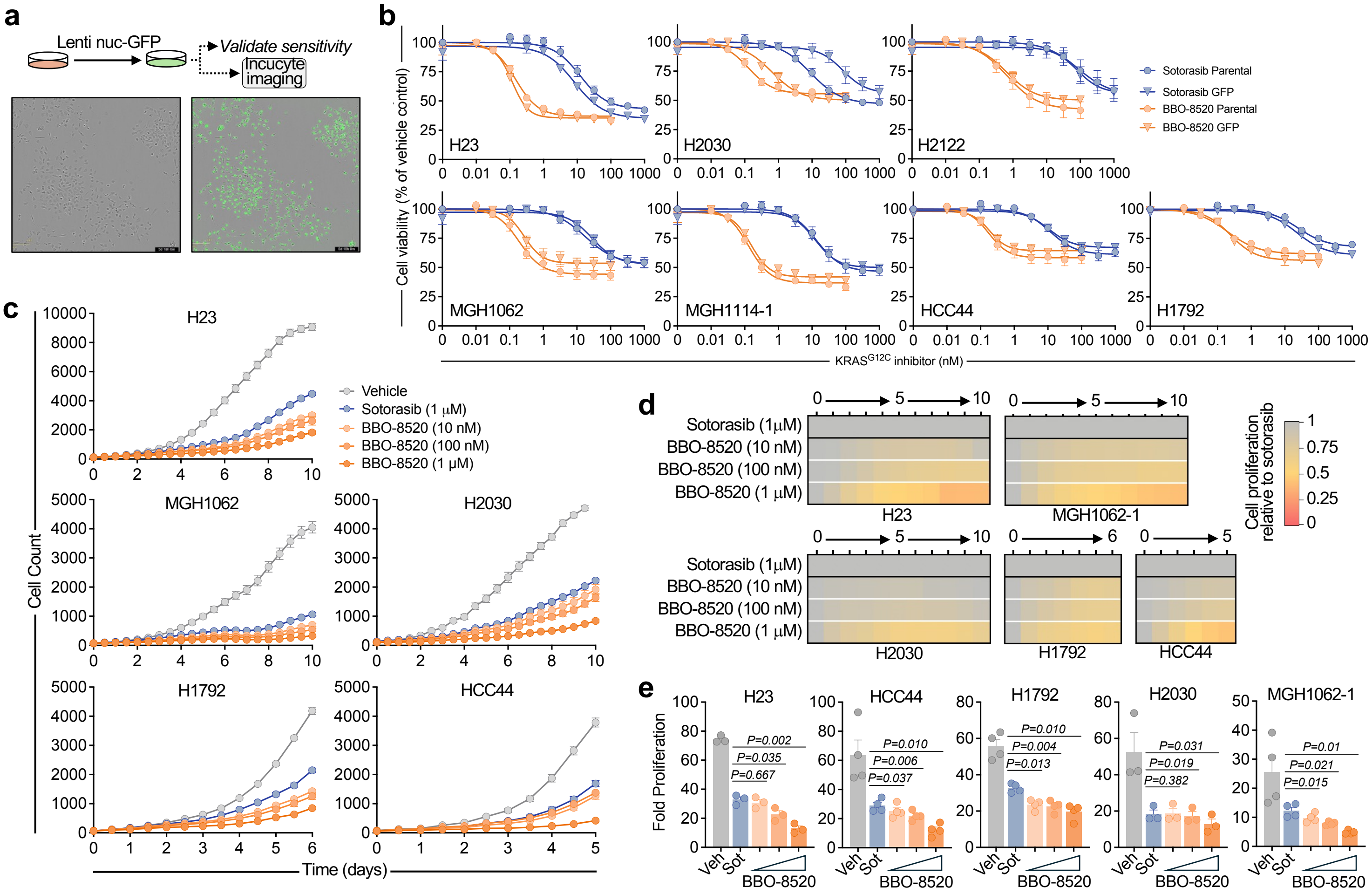


**Supplemental Figure 2. BBO-8520 exhibits more durable suppression of cell proliferation compared to sotorasib. a,** Generation of GFP-labeled cell lines for live cell imaging. Representative images of the H23 cell line before and after labeling are shown. **b,** Composite dose response curves of matched parental and GFP-labeled cell lines treated with KRAS inhibitors for 72 hours. Cell viability was quantified by CellTiter-Glo. GFP-labelled curves shown are mean of n=2 independent biological replicates. **c,** Imaged-based monitoring (Incucyte) of proliferation in cell lines treated with sotorasib or BBO-8520. Data are mean and S.E.M of n=4 technical replicates and are representative of n=3-4 independent biological replicates. **d,** Relative cell proliferation of cells treated with BBO-8520, normalized to sotorasib. Values are mean of n=3-4 biological replicates, corresponding to the cell counts shown in panel C. **e,** Comparison of cell proliferation after treatment with 10 nM, 100 nM or 1 μM BBO-8520 or 1 μM sotorasib. Values represent cell proliferation relative to baseline determined after 10 days (H23, MGH1062-1, H2030), 6 days (H1792) or 5 days (HCC44), from 3-4 independent biological replicates.

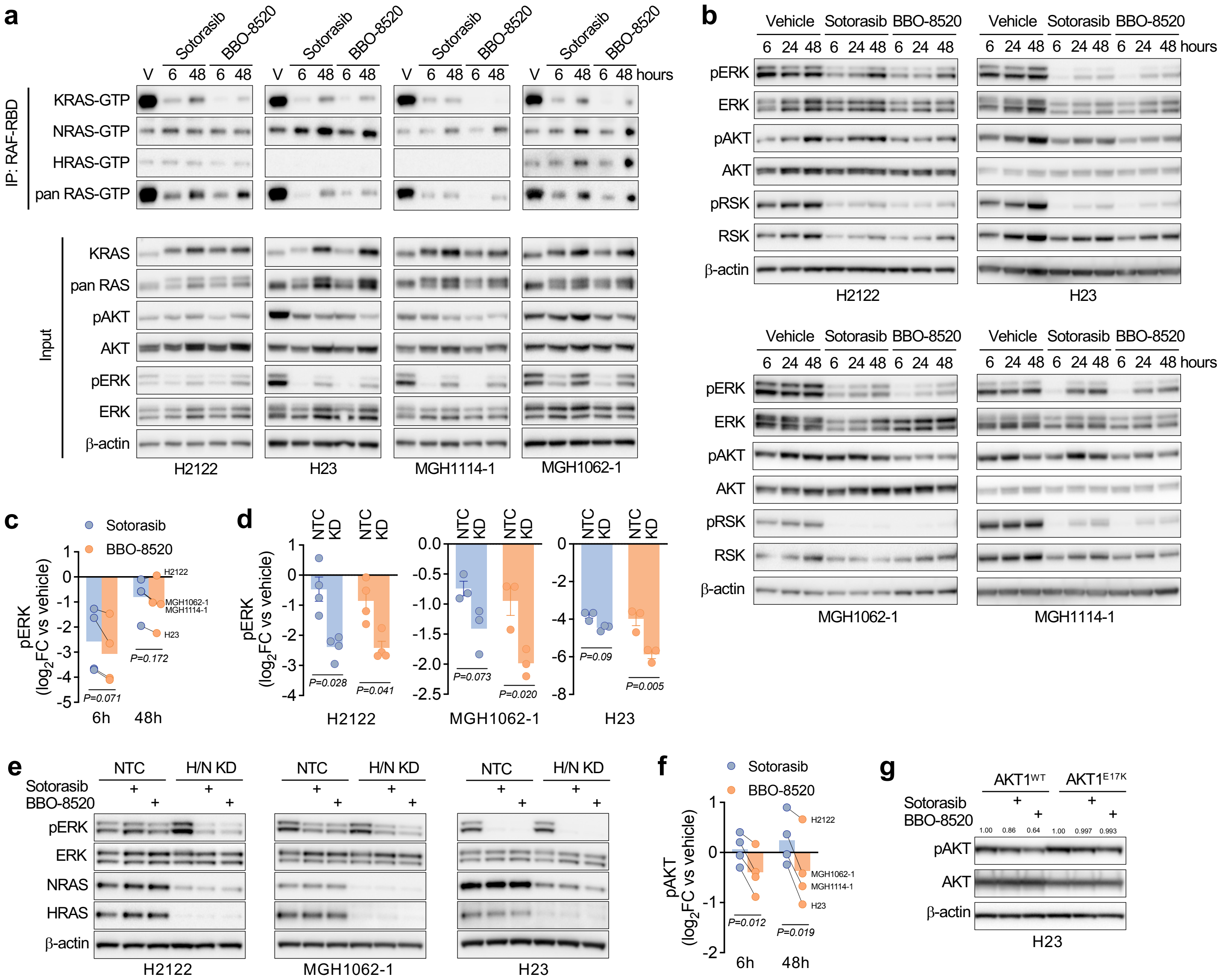


**Supplemental Figure 3. BBO-8520 achieves more durable suppression of KRAS$^{G12C}$-RAF engagement and inhibition of PI3K-AKT compared to sotorasib. a,** *KRAS$^{G12C}$*-mutant NSCLC cells were treated with BBO-8520 (100 nM) or sotorasib (1 µM) for 6 or 48 hours and RAF-RBD pull-down was performed to assess engagement between KRAS-GTP, NRAS-GTP, HRAS-GTP and RAF. Data are representative of n=3 independent biological replicates and correspond to quantified values in Figure 3A. **b,** Representative western blot images of cells treated with BBO-8520 (100 nM) or sotorasib (1 µM) for 6, 24, or 48 hours. Data are representative of n=7-8 independent biological replicates and correspond to the quantified values in panels C and F. **c,** Average change in phospho-ERK levels after treatment with sotorasib (1 µM) or BBO-8520 (100 nM) for 6 or 48 hours. Data are quantified band intensities from western blots (see panel B) normalized to vehicle control, n=7-8 independent biological replicates Each data point represents the average change in phospho-ERK in an individual cell line. **d-e,** Cell lines with siRNA knockdown of HRAS/NRAS (KD) or non-targeting control (NTC) were treated with sotorasib (1 µM) or BBO-8520 (100nM) for 48 hours and harvested for western blotting. Data in panel D are quantified band intensities normalized to vehicle treated cells, n=3 independent biological replicates. Panel E shows representative western blot of n=3 biological replicates. **f,** Average change in phospho-AKT (S473) levels after treatment with sotorasib (1 µM) or BBO-8520 (100 nM) for 6 or 48 hours. Data are quantified band intensities from western blots (see panel B) normalized to vehicle control, n=7-8 independent biological replicates Each data point represents the average change in phospho-AKT in an individual cell line. **g,** Western blot images of H23-AKT1$^{wt}$ and H23-AKT1$^{E17K}$ cells treated with BBO-8520 (100 nM) or sotorasib (1 µM) for 6 hours. Quantified band intensities of phospho-AKT (normalized to vehicle controls) are listed above the corresponding band.

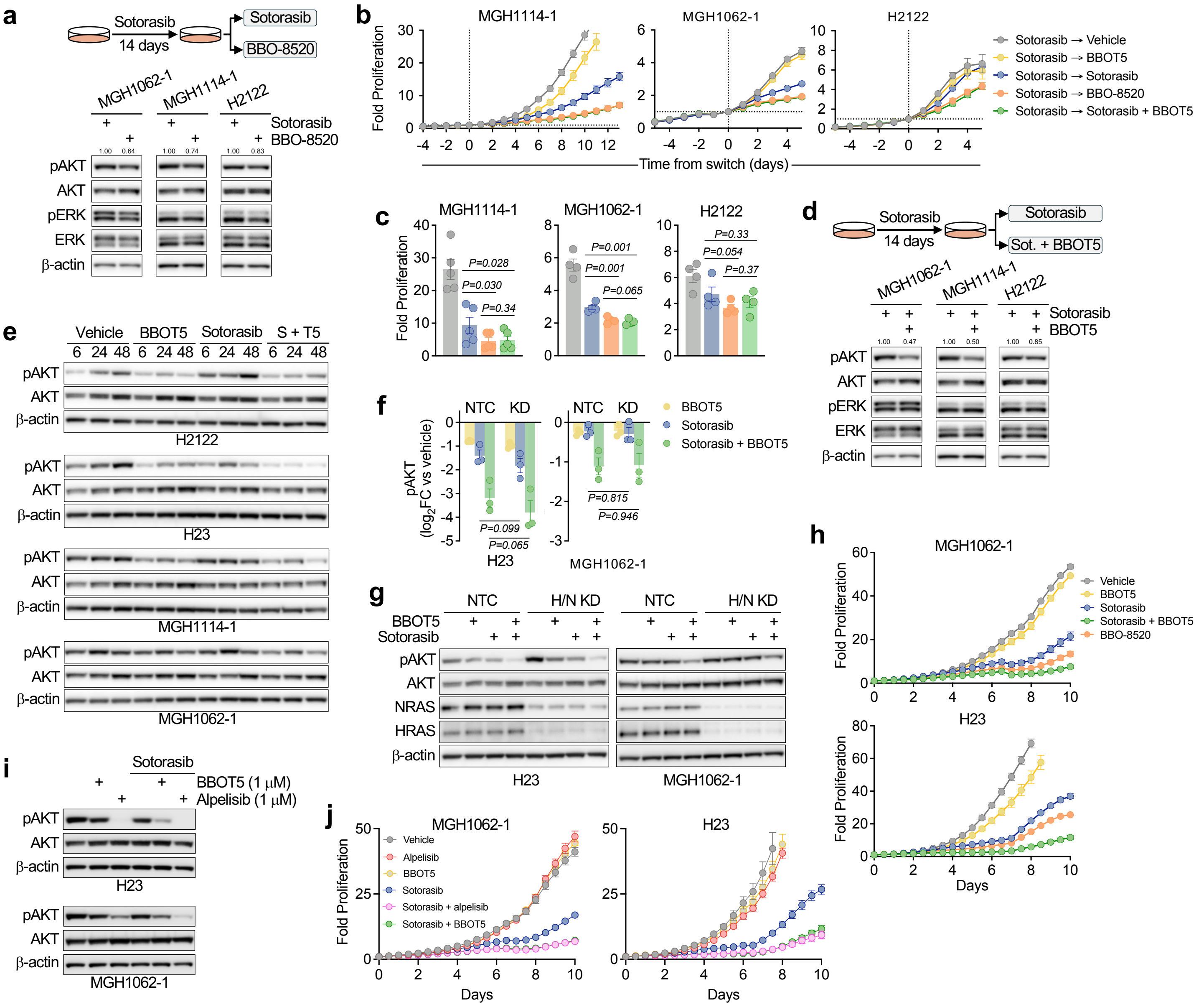


**Supplemental Figure 4. Disruption of RAS-PI3Kα increases sensitivity to sotorasib, phenocopying BBO-8520. a,** Cells were treated with sotorasib (1 μM) for 14 days, followed by switch to BBO-8520 (100 nM) or continued sotorasib for 24 hours, then harvested for western blot analysis. Quantified band intensities of phospho-AKT (normalized to sotorasib treatment) are listed above the corresponding band. **b,** Cell lines were treated with sotorasib (1 μM) for 7 days followed by switch to the indicated conditions (BBO-8520, 100 nM; BBOT5, 1 μM; sotorasib, 1 μM). Timepoints shown are relative to switch ("day 0"). Data are mean and S.E.M. of n=3-6 technical replicates and are representative of n=4 independent biological replicates. **c,** Comparison of cell proliferation after treatment as described in panel B. Data points represent mean cell proliferation 5 (MGH1062-1, H2122) or 13 (MGH1114-1) days after drug switch, normalized to the cell count at the time of drug switch, n=4 independent biological replicates. **d,** Cells were treated with sotorasib (1 μM) for 14 days, followed by addition of BBOT5 (1 μM) or continued sotorasib for 24 hours, then harvested for western blot analysis. Quantified band intensities of phospho-AKT (normalized to sotorasib treatment) are listed above the corresponding band. **e,** Cells were treated with BBOT5 (1 μM), sotorasib (1 μM), or combination for 6, 24, or 48 hours then harvested for western blot analysis. Data are representative of n=3 biological replicates and correspond to quantified values in Figure 4B. **f-g,** Cell lines with siRNA knockdown of HRAS/NRAS (KD) or non-targeting control (NTC) were treated with BBOT5 (1 μM), sotorasib (1 μM) or the combination for 48 hours and harvested for western blotting. Data in panel F are quantified band intensities normalized to vehicle treated cells, n=3 independent biological replicates. Panel G shows representative western blot of n=3 biological replicates. **h,** Cells were treated with sotorasib (1 μM), BBOT5 (1 μM), BBO-8520 (100 nM) or the indicated combination and cell numbers were quantified by live cell imaging (Incucyte). Data is mean and S.E.M. of n=3-4 technical replicates and is representative of n=3-4 independent biological replicates. **i,** Cell lines were treated with BBOT5 (1 μM), alpelisib (1 μM), sotorasib (1 μM) or combinations for 24 hours and harvested for western blotting. Data are representative of n=3-4 biological replicates. **j,** Cells were treated with sotorasib (1 μM), BBOT5 (1 μM), alpelisib (1 μM) or the indicated combinations and cell numbers were quantified by live cell imaging (Incucyte). Data is mean and S.E.M. of n=3 technical replicates and is representative of n=3 independent biological replicates.

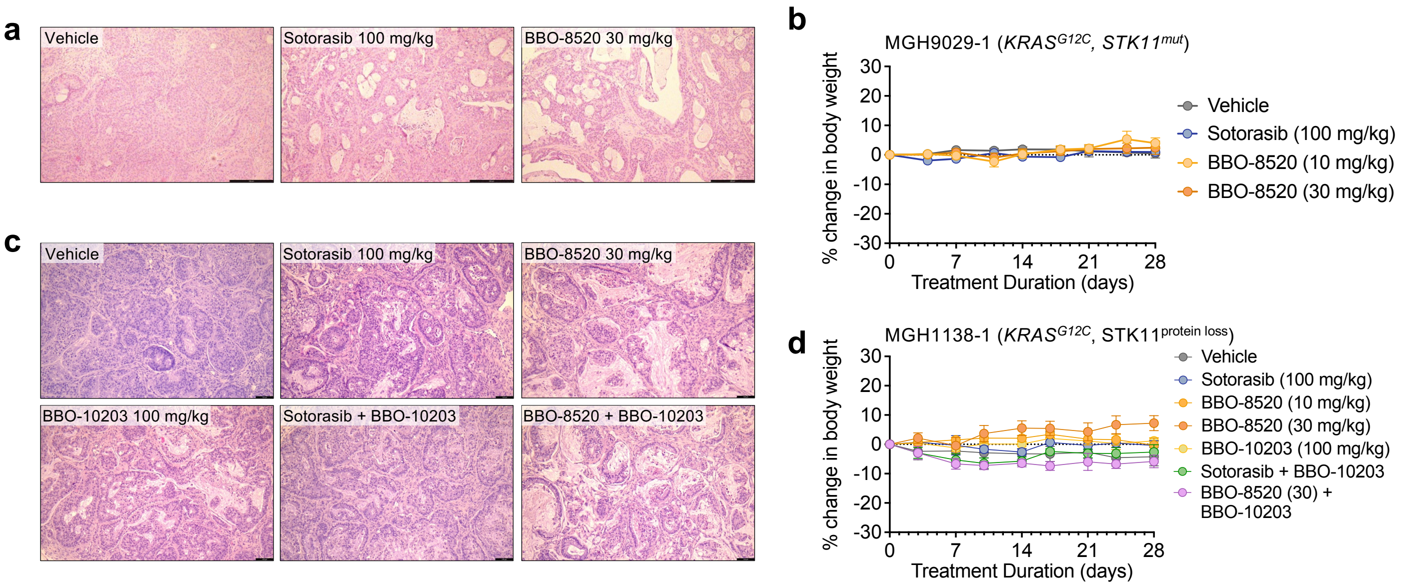


**e**

| Study Type | Time (hours) | [Sotorasib] (ng/mL (± SD)) 100 mg/kg |
|---|---|---|
| PK QDx1 (NSG mice) | 2 | 669 (+/- 64) |
| | 6 | 181 (+/- 74) |
| | 24 | 2 (+/- 1) |

| Study Type | Time (hours) | [BBO-8520] (ng/mL (+/- SD)) 10 mg/kg | 30 mg/kg |
|---|---|---|---|
| PK QDx1 (NSG mice) | 2 | 97 (± 32) | 348 (± 105) |
| | 6 | 92 (± 37) | 255 (± 72) |
| | 24 | 5 (± 3) | 29 (± 12) |

| | BBO-8520 (10 mg/kg) | BBO-8520 (30 mg/kg) | Sotorasib (100 mg/kg) |
|---|---|---|---|
| AUC 0-24 (h*ng/mL) | 1353 | 4139 | 4018 |
| fu | 0.0026 | 0.0026 | 0.06 |
| Free AUC 0-24 (h*ng/mL) | 3.5 | 10.8 | 241.1 |

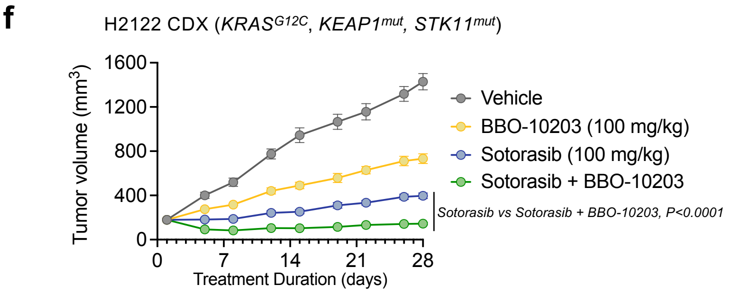


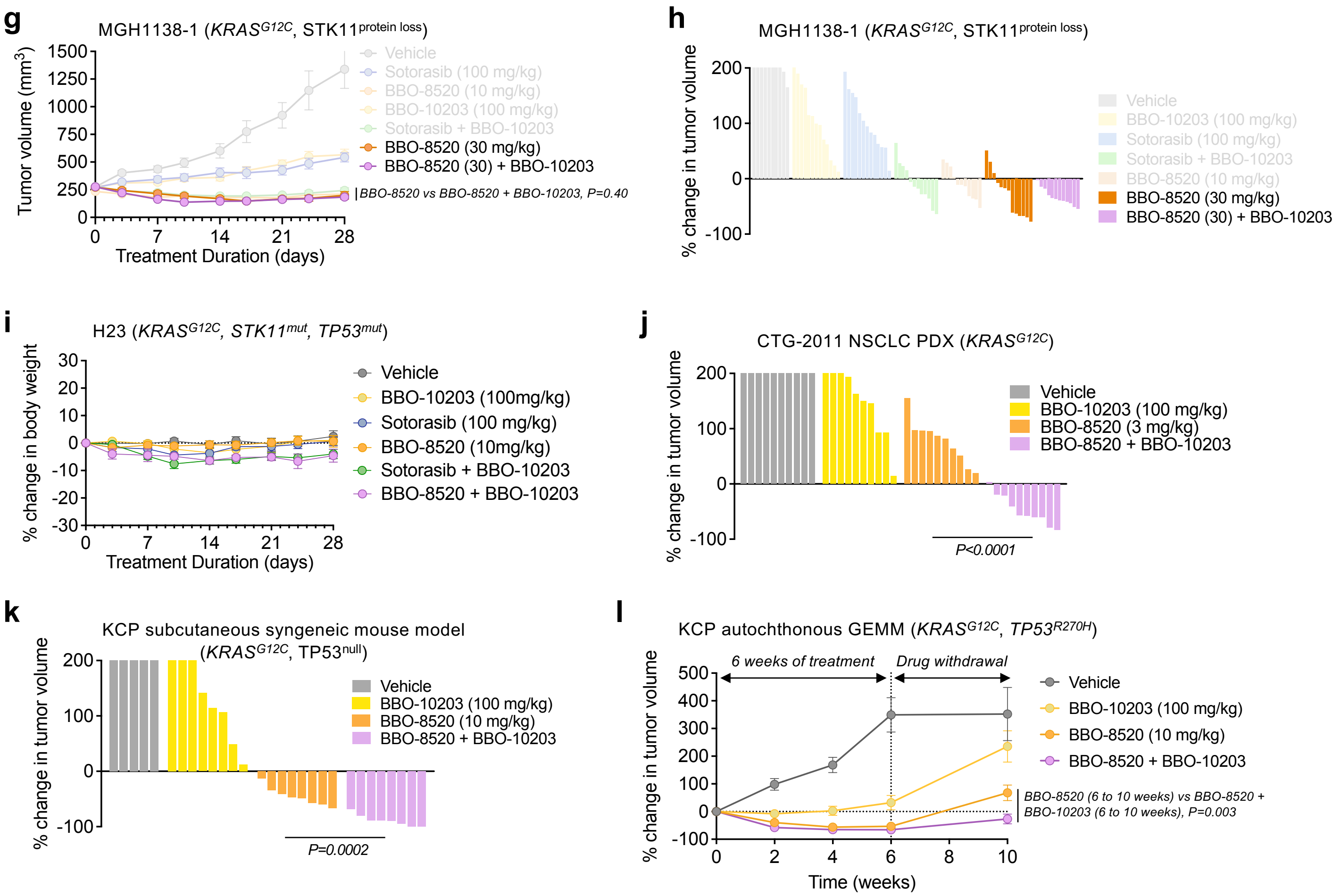


Supplemental Figure 5

**Supplemental Figure 5. Disruption of RAS-PI3Ka with BBO-10203 increases *in vivo* efficacy of KRAS (OFF) and KRAS (ON) inhibitors. a,** Representative H&E images of MGH9029-1 xenograft tumors after 3 days treatment. **b,** Change in body weight of mice bearing MGH9029-1 xenograft tumors during drug treatment. Data are mean and S.E.M. (vehicle, n=11; sotorasib100 mg/kg, n=12; BBO-8520 10 mg/kg, n=13; BBO-8520 30 mg/kg, n=15). **c,** Representative H&E images of MGH1138-1 xenograft tumors after 3 days treatment. **d,** Change in body weight of mice bearing MGH1138-1 xenograft tumors during drug treatment. Data are mean and S.E.M. (vehicle, n=10; sotorasib 100 mg/kg, n=13; BBO-8520 30 mg/kg, n=11; BBO-10203 100 mg/kg, n=13; sotorasib + BBO-10203, n=12). **e,** Pharmacokinetic parameters of BBO-8520 and sotorasib in NSG mice. **f,** Mice bearing H2122 xenograft tumors were treated with vehicle (n=10), sotorasib (100 mg/kg, n=10), BBO-10203 (100 mg/kg; n=10), or sotorasib + BBO-10203 (n=10) once daily by oral gavage. Data are mean and S.E.M. **g,** Mice bearing MGH1138-1 PDX tumors were treated with vehicle (n=10), sotorasib (100 mg/kg, n=13), BBO-8520 (30 mg/kg, n=11), BBO-10203 (100 mg/kg; n=13), or sotorasib + BBO-10203 (n=12) once daily by oral gavage. BBO-8520 30 mg/kg and BBO8520 30 mg/kg + BBO=10203 are shown; the other treatment arms are replotted from Figure 5C for comparison purposes. Data are mean and S.E.M. **h,** Waterfall plot of the change in tumor volume at 28 days compared to baseline. BBO-8520 30 mg/kg and BBO8520 30 mg/kg + BBO=10203 are shown; the other treatment arms are replotted from Figure 5D for comparison purposes. **i,** Change in body weight of mice bearing H23 xenograft tumors during drug treatment. Data are mean and S.E.M. (vehicle, n=8; sotorasib 100mg/kg, n=8; BBO-8520 10 mg/kg, n=11; BBO-10203 100 mg/kg, n=13; sotorasib + BBO-10203, n=8; BBO-8520 + BBO-10203, n=8. **j,** Waterfall plot of the change in tumor volume of CTG-2011 xenograft tumors at 28 days compared to baseline. **k,** Waterfall plot of the change in tumor volume of KCP tumors at 28 days compared to baseline. **l,** Mice bearing autochthonous KCP tumors were treated with vehicle (n=14), BBO-8520 (10 mg/kg, n=15), BBO-10203 (100 mg/kg, n=11), or BBO-8520 + BBO-10203 (n=13) once daily by oral gavage for 6 weeks followed by 4 weeks with no treatment. Data are mean and S.E.M.
